# Microfluidic immuno-serology assay revealed a limited diversity of protection against COVID-19 in patients with altered immunity

**DOI:** 10.1101/2022.08.31.506117

**Authors:** Dongjoo Kim, Giulia Biancon, Zhiliang Bai, Jennifer VanOudenhove, Yuxin Liu, Shalin Kothari, Lohith Gowda, Jennifer M. Kwan, Nicholas Carlos Buitrago-Pocasangre, Nikhil Lele, Hiromitsu Asashima, Michael K. Racke, JoDell E. Wilson, Tara S. Givens, Mary M. Tomayko, Wade L. Schulz, Erin E. Longbrake, David A. Hafler, Stephanie Halene, Rong Fan

## Abstract

The immune response to SARS-CoV-2 for patients with altered immunity such as hematologic malignancies and autoimmune disease may differ substantially from that in general population. These patients remain at high risk despite wide-spread adoption of vaccination. It is critical to examine the differences at the systems level between the general population and the patients with altered immunity in terms of immunologic and serological responses to COVID-19 infection and vaccination. Here, we developed a novel microfluidic chip for high-plex immuno-serological assay to simultaneously measure up to 50 plasma or serum samples for up to 50 soluble markers including 35 plasma proteins, 11 anti-spike/RBD IgG antibodies spanning all major variants, and controls. Our assay demonstrated the quintuplicate test in a single run with high throughput, low sample volume input, high reproducibility and high accuracy. It was applied to the measurement of 1,012 blood samples including in-depth analysis of sera from 127 patients and 21 healthy donors over multiple time points, either with acute COVID infection or vaccination. The protein association matrix analysis revealed distinct immune mediator protein modules that exhibited a reduced degree of diversity in protein-protein cooperation in patients with hematologic malignancies and patients with autoimmune disorders receiving B cell depletion therapy. Serological analysis identified that COVID infected patients with hematologic malignancies display impaired anti-RBD antibody response despite high level of anti-spike IgG, which could be associated with limited clonotype diversity and functional deficiency in B cells and was further confirmed by single-cell BCR and transcriptome sequencing. These findings underscore the importance to individualize immunization strategy for these high-risk patients and provide an informative tool to monitor their responses at the systems level.

## Main

Coronavirus disease-19 (COVID-19) has become a serious worldwide public health emergency and the ongoing evolution of the SARS-CoV-2 virus still poses enormous challenges for global pandemic control^1^. Compared to healthy subjects, patients with hematologic malignancies or autoimmune disease tend to suffer more severe and prolonged courses of COVID-19 infection and are at higher risk of developing severe acute respiratory syndrome, due largely to their altered immune fitness condition such as myelosuppression and lymphopenia^2-4^. Although vaccinations are highly effective against symptomatic disease and notably reduce fatality rates^5^, patients with hematologic malignancies such as chronic lymphocytic leukemia (CLL), multiple myeloma (MM), and autoimmune diseases under immunosuppressive treatments may not mount adequate neutralizing antibody responses after receiving vaccinations^6,7^. Thus, as COVID-19 variants continuously emerge, there is still an unmet need for new immuno-serological assays to systematically evaluate the effectiveness of immune protection against COVID-19 in these vulnerable populations.

Microfluidic chips are a technological platform of choice for performing a rapid test of plasma protein biomarkers including SARS-CoV-2 specific serum antibodies in conjunction with the markers of immunocompetence in a high-throughput, high sensitivity and specificity, low blood sample consumption, and low-cost manner. Several new technologies have been developed and have demonstrated superior detection performances than traditional methods such as enzyme-linked immunosorbent assays^8^, chemiluminescent immunoassays^9^ and lateral flow assays^10^. For example, Swank and colleagues developed and validated a nanoimmunoassay device to detect the presence of anti-spike IgG antibodies with throughput of 512 to 1,024 samples in parallel^11^, and Rodriguez-Moncayo et al. reported a 4-plex SARS-CoV-2 serology platform that can measure 50 serum samples per assay^12^. These platforms routinely show high detection sensitivity (>95%) and specificity (>90%) with ultralow-volume whole blood or plasma samples as input (less than 2 μL). However, due to the limitation of current microfluidic pattern design, the level of multiplexing in these devices remains low (less than 10) and most assays are limited to the detection of only antibody responses. As new SARS-CoV-2 variants continue to emerge, there is a constant demand for assays that offer high-plex co-profiling of antibody binding against all variants of concern. Moreover, considering the orchestrated immune responses induced by infection or vaccination, a device capable of simultaneously detecting antibodies and immune mediators would be highly desirable.

Here we report a portable microdevice designed to perform highly multiplexed measurements of anti-SAR-CoV-2 antibodies and immune mediator proteins (up to 50 in total) in microliters of human serum with high sample throughput of up to 50 samples per run per device. A “microfluidic patterning chip” was prepared in advance to perform microchannel-guided immobilization of capture antibodies or viral antigens, creating a 1D protein stripe barcode array on a glass slide^13^. When patient serum samples are ready to measure, a “microfluidic test chip” with a set of parallel microchannels allows for loading up to 50 serum samples over the microarray for simultaneous detection of serum proteins and SARS-CoV-2 IgG antibodies. We successfully co-measured 48 human serum samples plus one labeling control to detect a panel of 35 soluble proteins and 11 anti-SARS-CoV-2 IgG proteins in a single assay. A total of 1,012 serum samples were successfully measured, demonstrating high robustness of our high-plex immuno-serology assay. In this work, we performed in-depth analysis using 366 samples covering 127 patients with hematologic or other types of cancers, autoimmune disorders either with SAR-CoV-2 infection or pre- or post-COVID vaccination, and 21 healthy donors pre- or post-vaccination. We observed substantial differences of anti-spike/RBD IgG binding antibody (bAb) responses and immune context between patients with hematologic malignancies or receiving treatments compromising immune status and Non-Heme Cancer patients and healthy controls in response to both COVID infection and vaccination. As expected, our assay revealed limited IgG bAb response in those with a weakened immune system as compared to healthy donors and provided a comprehensive assessment of multi-factorial immune responses. We also found an impaired RBD-specific binding ability in hematologic malignancy patients with COVID infection that could be associated with a limited diversity of B cell clonotypes, which was confirmed by single-cell B cell repertoires (BCR) sequencing.

## Results

### Development and calibration of high-plex immuno-serology assay

To develop high-plex immuno-serology assay with high versatility and low complexity, we utilized a single-layer microfluidic design strategy that employs a polydimethylsiloxane (PDMS) “microfluidic patterning chip” and a poly-L-lysine coated glass slide (PLL slide) for creating high-density protein barcode array and a “PDMS microfluidic test chip” for multiplexed measurement of protein targets from a large number of samples (Figure 1a). This microfluidic flow patterning method as described in our previous work^13^ was adapted to fabricate high-density barcode array chips. In total, 35 capture antibodies, including antibodies against cytokines/chemokines, angiogenesis markers, neutrophil activation markers, endotheliopathy markers, and 11 SARS-CoV-2 recombinant antigens were immobilized on the PLL slide by using the first PDMS chip, which contains 50 parallel microfluidic channels with 5-turn serpentine patterns in each channel (Figure 1b). After removing the first PDMS chip, the immobilized functional microarray slide can be stored in -80°C freezer until use. When serum samples are ready to measure, a second PDMS test chip with 50 microchannels perpendicular to those in the first PDMS chip was placed on the microarray slide. This unique design allows simultaneous measurement of 50 serum samples with 5 replicates for each sample, minimizing the detection error and/or bias caused by nonspecific antibody bindings or debris (Figure 1c). After flowing the serum samples over the surface of the microarray, the second microfluidic chip was removed and a cocktail mixture of biotinylated detection antibodies and fluorescent dye conjugated anti-human IgG antibodies was loaded onto the microarray slide. Finally, the fluorescent images were obtained by using a three-laser microarray scanner (Genepix 4200) and a software suite has been developed to quantify fluorescence intensities of the corresponding protein targets (Methods).

**Figure 1.**
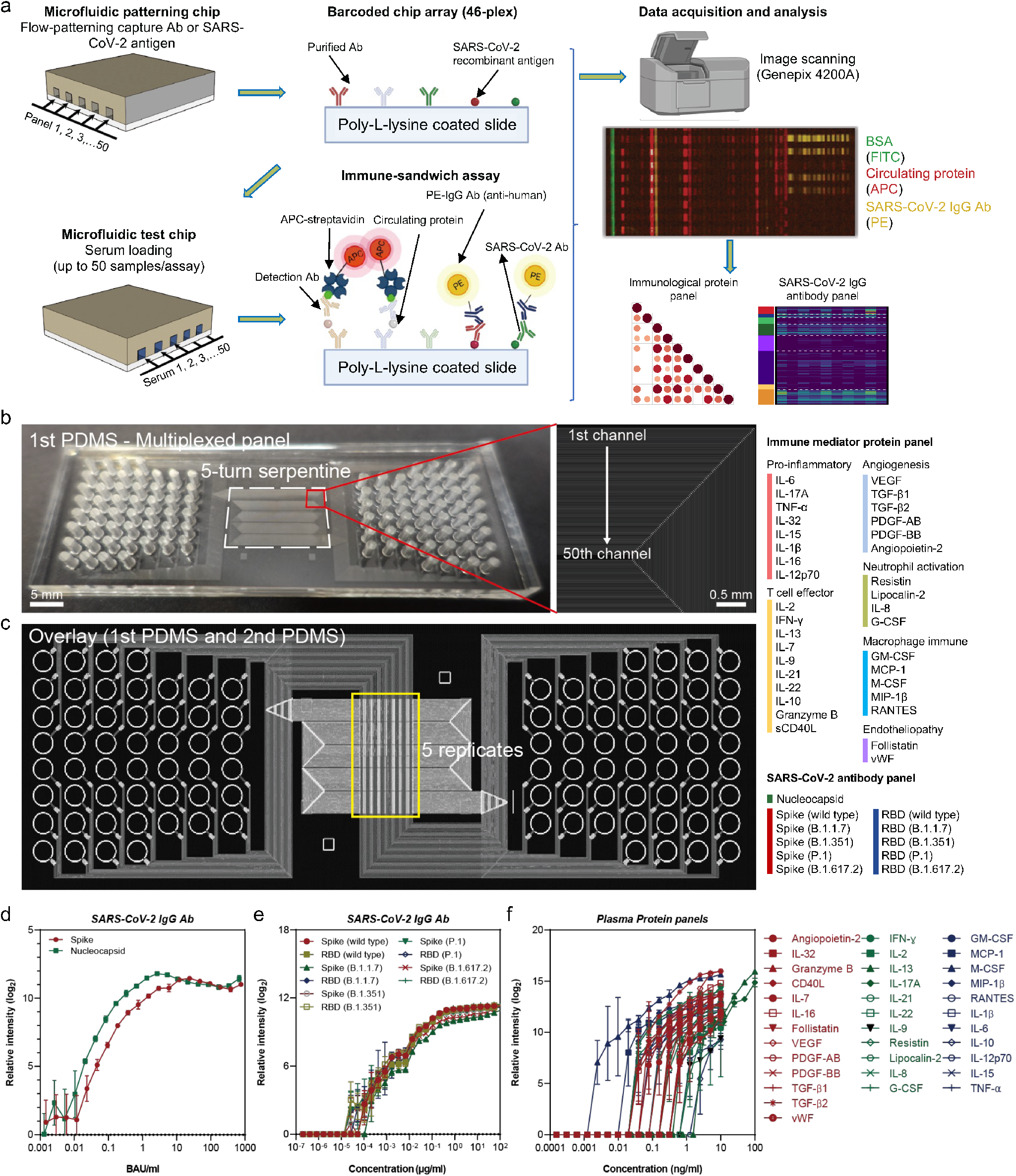
High-plex immuno-serology assay design and titration test. **(a)** Schematic workflow. The first PDMS “microfluidic patterning chip” with 50 parallel microchannels was placed on the poly-l-lysine coated glass slide (PLL slide), then purified antibodies or SARS-CoV-2 recombinant antigens were flowed into the microchannels. After removing the first PDMS, the second “microfluidic test chip” was placed on the same PLL slide. Then, serum samples were added, and the captured proteins or SARS-Cov-2 binding antibodies were detected via a surface-bound immune-sandwich assay. Finally, scanning fluorescent images were obtained and analyzed. **(b)** Photographic image of the first PDMS microchip. **(c)** Integrated device design after overlaying the first and second PDMS. The protein and SARS-CoV-2 serology panels evaluated in this work are listed on the right side. **(d)** The titration curves of US SARS-CoV-2 serology standard (Frederick National Laboratory). **(e)** The titration curves of SARS-CoV-2 anti-spike/RBD antibodies. **(f)** The titration curves of protein panels. Each titration curve was plotted with hyperbola equation in nonlinear regression. The data presented is the mean value of 5 replicates from a single assay. Scatter plots show means ± SEM. PDMS, polydimethylsiloxane; Ab, antibody; BSA, bovine serum albumin; FITC, fluorescein isothiocyanate; APC, allophycocyanin; PE, phycoerythrin; BAU, Binding Antibody Units.

To quantitatively evaluate the detection ability of our device, we conducted titration tests using recombinant proteins, anti-SARS-CoV-2 Spike Abs, and the US SARS-CoV-2 serology standards. Serially (2-fold) diluted antigen or SARS-CoV-2 Ab solutions in 1X phosphate-buffered saline (PBS) were loaded in each inlet of the microfluidic test device and flowed onto the barcode array. The relative intensity against recombinant proteins or SARS-CoV-2 Spike Ab was adjusted by log2 normalization after subtracting the background threshold. We first utilized Human SARS-CoV-2 Serology Standard Spike IgG and Nucleocapsid IgG generated by the NCI Frederick National Laboratory for Cancer Research (FNLCR) to evaluate the ligand binding activity of our assay, demonstrating a detection range of 0.01~1000 Binding Antibody Units (BAU)/ml (Figure 1d). Then, commercially available SARS-CoV-2 Abs were used to estimate the concentrations of IgG Ab against spike or RBD antigens of SARS-CoV-2 wild type and other variants, including alpha (B.1.1.7), beta (B.1.351), gamma (P.1), and delta (B.1.617.2). We found that titration curves from all recombinant antigens merged together with notably high Pearson’s R values (Figure 1e and Figure S1), indicating comparable binding affinities of commercial antibodies to the panel of antigens. There are no standard values available in these antibodies to convert the fluorescent intensities of theses variants to BAU/ml. However, we were able to choose the titration curve of wild type spike antigen as a reference and convert all the other scanning intensities to concentrations. A detection capacity ranging from 0.001μg/ml to 100μg/ml was achieved for these IgG bAbs (Figure 1e). Similarly, the titration curve for a 35-protein panel was generated and a dynamic range of 0.001~10 ng/ml was obtained (Figure 1f). Overall, these data indicate that our high-plex immuno-serology assay could quantify concentrations of both SARS-CoV-2 antibodies and plasma protein markers with good sensitivity and dynamic range.

### Validation of assay performance

We first demonstrate the throughput capacity of our high-plex assay with 48 serum samples obtained from patients. After fabricating a microarray by immobilizing SARS-CoV-2 recombinant antigens and capture antibodies against a panel of immune mediator proteins, 8-μl volumes of serum sample from each patient/donor were loaded onto the device and drawn from the inlets to the outlets through the barcode array for 1 hour using a controlled house vacuum system. To differentiate the location of the starting column and row in the raw fluorescent images, we introduced the fluorescein isothiocyanate (FITC)-conjugated bovine serum albumin (BSA) and FITC-conjugated anti-mouse IgG Ab to the first channel of two PDMS microfluidic chips, respectively. Allophycocyanin (APC, 635 nm emission) and phycoerythrin (PE, 532 nm emission) were employed to measure circulating plasma proteins and anti-SARS-CoV-2 bAbs, respectively. In the scanned fluorescent image of the assay region (Figure 2a), each row represents one tested sample and each column is the signal of a specific protein or anti-SARS-CoV-2 antibody. We observed high consistency of the florescence patterns between the 5 replicates among all the 48 samples, indicating the robustness of our assay. For data quantification, we exported the fluorescence intensity and converted it to concentration or BAU/ml using the titration curves (Figure S2a,b).

**Figure 2.**
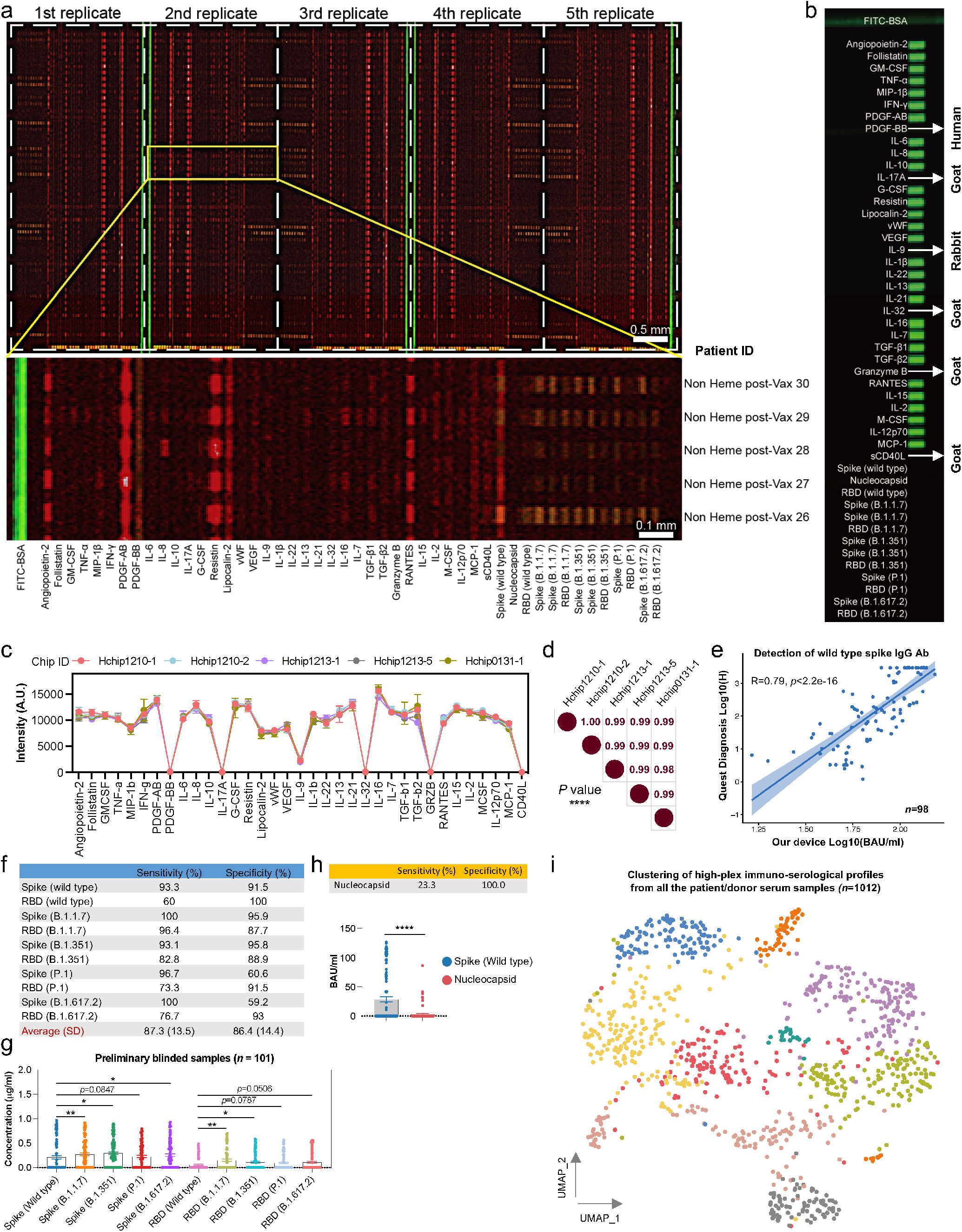
Validation of assay performance. **(a)** Representative fluorescent image of high-plex immune-serology assay in the measurement of 48 serum samples in a single run. Red and yellow signal represent the plasma protein and SARS-CoV-2 IgG Ab, respectively. Enlarged image shows a fluorescence signal from 5 serum samples. FITC conjugated BSA was introduced to differentiate first column. **(b)** The evaluation of potential channel cross-talk. Antibodies generated from other species were coated for the indicated proteins, and the remaining 29 are mouse-derived capture antibodies. FITC-conjugated anti-mouse IgG Ab was introduced to the first row. **(c)** Device-to-device reproducibility evaluation using 5 barcoded array chips prepared at different time points. **(d)** The Pearson correlations across all the 5 chips. **(e)** The correlation plot between our high-plex assay and commercialized Quest diagnosis in the measurement of 98 serum samples. **(f)** The sensitivity and specificity of each evaluated anti-spike/RBD antibody. 101 preliminary blinded samples from NCI-Frederick National Laboratory for Cancer Research were used. **(g)** The concentration comparisons between anti-spike or anti-RBD antibodies. **(h)** The sensitivity and specificity of nucleocapsid in the test of 101 preliminary blinded samples. **(i)** Unsupervised clustering of all the 1012 samples measured using our assay (only for visualization). The concentrations of the 35 proteins and 10 anti-spike/RBD antibodies were used to perform the clustering. Each point represents one measured sample. *P* values were calculated with two-tailed Mann-Whitney test. (* *P* < 0.05, ** *P* < 0.01, *** *P* < 0.001, **** *P* < 0.0001). Scatter plots show means ± SEM. Ab, antibody; BSA, bovine serum albumin; FITC, fluorescein isothiocyanate.

To confirm that there is no crosstalk or leakage between channels during the reagent flowing, we coated antibodies generated from other species (human, goat or rabbit) for PDGF-BB, IL-17A, IL-9, IL-32, Granzyme B and sCD40L, whereas the remaining 29 are mouse-derived capture antibodies. After applying FITC-conjugated anti-mouse IgG Ab to the first row of the second PDMS device, no signal was detected for the above-mentioned proteins as well as the SARS-CoV-2 spike or RBD antigens, suggesting minimum crosstalk (Figure 2b). To assess device-to-device reproducibility, we repeated the tests independently using 5 barcoded array chips fabricated at different times (Figure 2c), and the results show tight correlation of average Pearson’s R=0.99 (*p* value <0.0001) (Figure 2d).

To explore the accuracy and reliability of our device, we compared our high-plex serology assay with a commercially available assay from Quest Diagnostics. Our multiplex assay had a Pearson’s R of 0.79 (*p* value <0.0001) when correlated with the commercial, semiquantitative assay for anti-SARS-CoV-2 Spike IgG bAbs for 98 serum samples (Figure 2e). We then evaluated all the anti-spike and RBD antibodies in our serology panel using 101 blinded samples provided by NCI FNLCR including sera from patients with COVID infection and healthy controls (Figure S3a-c). Independent analysis by scientists at the NCI FNLCR revealed that overall average sensitivity and specificity of IgG reactivity are 87.3% and 86.4%, respectively (Figure 2f). Specifically, the detection sensitivity of IgG against spike antigens was higher than against the RBD (mean 96.6% vs. mean 77.8%), whereas the specificity of RBD detection was greater in comparison with that of spike antibodies (mean 92.2% vs. mean 80.6%). The best overall performance was obtained by the detection of IgG against alpha (B.1.1.7) spike with 100% sensitivity and 95.9%, specificity. Moreover, for both anti-spike and RBD antibodies, we found that later-stage variants were generally detected at higher sensitivity and lower specificity than wild type. Nevertheless, concentration levels were not affected by differences in sensitivity/specificity (Figure 2g). Consistent with this observation, we found that the antibody binding affinities for the nucleocapsid protein were significantly lower than for wild type Spike antigen, suggesting sub-optimal performance of this assay in the current configuration to distinguish vaccination from natural infection (Figure S3d).

We further conducted a comparative study to validate the detection performance of the immune mediator protein panel. Serum samples from four healthy controls, four intensive care unit (ICU)-survived patients, and four deceased patients were obtained and evaluated using our high-plex device. Although few significant differences were observed due to the limited cohort size, our data showed a clear trend towards elevated concentrations of IL-8, GM-CSF, resistin, G-CSF, IFN-γ, IL-6, IL-17A, lipocalin-2, MIP-1β, and TNF-α in ICU-survived or ICU-deceased patients compared with controls (Figure S4a), which is highly consistent with the results from a third party commercial assay in a previously published study that profiled these proteins in the same patient populations (Figure S4b)^14^. Thus, all these fully validated our microfluidic immune-serology assay using the commercial serology test by Quest Diagnostics, the blind samples from NCI FNLCR, and the commercial multiplex protein assay in the same patient cohort. Our device can perform 46-plex detection of SARS-CoV-2 bAbs and functional proteins at a throughput of up to 50 serum samples in a single run, with low sample volume input, minimal crosstalk, high reproducibility and high accuracy. Notably, this high-plex assay has been used to measure 1,012 serum samples to monitor immune responses of natural infection or vaccination and this cohort is still expanding (Figure 2h). The highly consistent measurements from standard samples and the real world clinical serum samples have demonstrated the robustness of our microfluidic high-plex immune-serology assay (Figure S5).

### Association of immune mediator proteins in 148 patients and donors upon vaccination

We conducted in-depth assessment in 148 patients or donors using our high-plex immuno-serology assay. Patients were divided into six groups based on their clinical demographics, immune status or baseline illness, and vaccination record (Figure S6a). Within our cohort, 38 patients were infected with acute COVID-19, 26 of them were patients with hematologic malignancies (Heme COVID+), and the remaining 12 were patients with non-hematologic cancers (Non-Heme Cancer COVID+). An additional 89 patients donated blood pre- and post-vaccination. Of these 23 had cancer and were known not to have had COVID infection prior to their blood draw (Non-Heme Cancer pre-Vax and Non-Heme Cancer post-Vax), and 66 patients had autoimmune diseases for which they received B cell depletion therapy (Autoimmune B cell depleted pre-Vax and Autoimmune B cell depleted post-Vax). Additionally, we included 21 healthy donors pre- and post-vaccination (HD pre-Vax and HD post-Vax) to establish baseline controls. Basic demographic information of these patients and donors is provided in Table S1. In regard to vaccine type, BNT162b2 (Pfizer-BioNTech, Germany), mRNA-1273 vaccine (Moderna, USA), or Ad26.COV2.S (Johnson & Johnson, USA) were administrated to patients/donors. In total, 366 serum samples were collected and analyzed.

Among all 35 immune mediators, we observed higher concentrations of PDGF-AB, IL-10, IL-12p70, and IL-13 compared to other protein markers (Figure S6b). To gain insights into key differences in these immune function proteins between patients with and without COVID infection and pre-versus post-vaccination, we compared the measurements of the 35 proteins across all groups and identified substantial differences in expression patterns. In contrast to minimal associations in healthy donors before vaccination (Figure S7a), we found several subsets of proteins whose expression levels are highly correlated in the patient groups. Specifically, in the Non-Heme Cancer Pre-Vax group, positive associations between T cell functional markers, including GM-CSF, TNF-α, IL-6, IL-16, IL7, and IL-15, dominated the correlation pattern (Figure 3a); patients with auto-immune diseases who received B-cell depletion therapy showed concerted dysregulation in a larger number of proteins in our panel albeit with lower correlation coefficients (Figure 3b). This suggests that our device and our protein panel can capture highly relevant protein signatures and associations unique to the responses of these vulnerable patient populations. Interestingly, we found a negative correlation between Resistin (RETN) and other markers of neutrophil activation including G-CSF and Lipocalin-2, and similar anticorrelation was also present in two angiogenesis signatures (PDGF-AB and PDGF-BB), implying complex immune dysregulation in these immunocompromised patients. After COVID vaccination, a concerted immune response emerged in healthy donors (HD post-Vax group) (Figure S7b), indicating that our device can detect systemic modulation of the immune responses post-vaccination. In Non-Heme Cancer patients, compared to HD, we found more modest changes in protein associations between pre- and post-vaccination samples (Figure 3c). Finally, as would be expected in immunocompromised patients, immunization did not induce coordinated immunological responses in Autoimmune (B cell depleted) patients as reflected in an unchanged correlation pattern in the post-Vax compared to pre-Vax setting in this patient population (Figure 3d).

**Figure 3.**
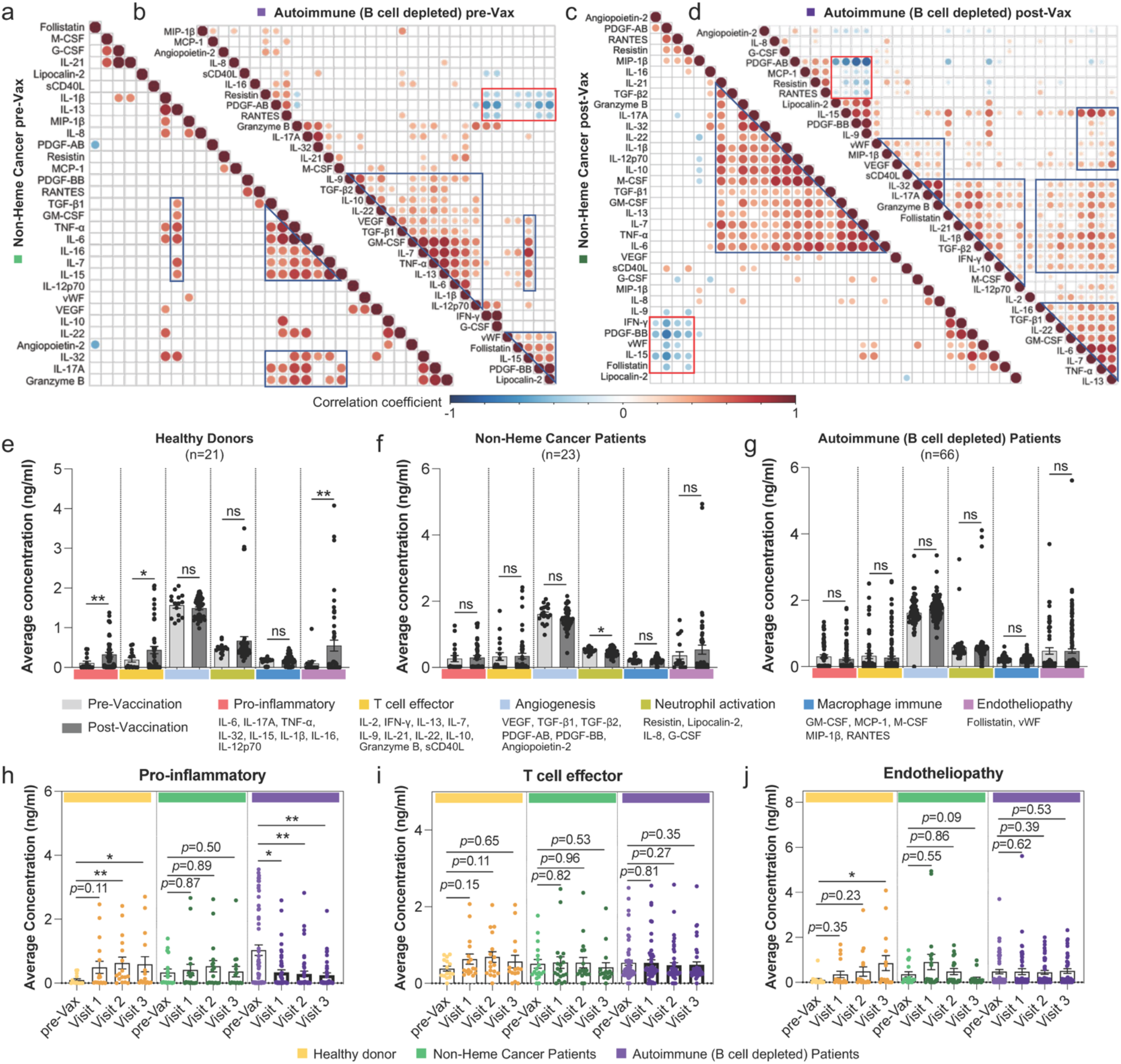
Vaccination-induced immunological functional protein response in each patient and donor group. **(a-d)** Correlation matrices of the 35 proteins evaluated in our high-plex immuno-serology assay panel for Non-Heme cancer patients pre-vaccination (a) and post-vaccination (c); autoimmune patients with B cell depletion therapy pre-vaccination (b) and post-vaccination (d). Only significant correlations (<0.05) are represented as dots. Pearson’s correlation coefficients from comparisons of protein concentrations across all the patients in a specific group are visualized by color intensity. Proteins were ordered by hierarchical clustering. **(e-g)** Comparisons of the average concentration of each functional protein category between pre- and post-vaccination in healthy donors (e); Non-Heme cancer patients (f); and autoimmune (B cell depleted) patients (g). **(h-j)** Comparisons of the average concentration of proteins regulating pro-inflammatory pathways (h); effector T cells (i); and endotheliopathy (j) between pre-vaccination and different timepoints post-vaccination. Visit 1, two weeks after the first vaccine dose; Visit 2, 0~3 days before the second vaccine dose; Visit 3, two weeks after the second vaccine dose. *P* values were calculated with two-tailed Mann-Whitney test. (* *P* < 0.05, ** *P* < 0.01, *** *P* < 0.001, **** *P* < 0.0001). ns, not significant.

We next sought to quantitively compare functional protein levels pre- and post-vaccination. As expected, in the healthy donor group, post-vaccination samples showed significantly higher concentrations of soluble proteins related to pro-inflammatory, T cell effector and endotheliopathy pathways, whereas proteins that regulate angiogenesis, neutrophil activation and macrophages remained mostly unchanged compared to pre-vaccination samples (Figure 3e). Non-Heme Cancer patients and patients with autoimmune disorders receiving B-cell depleting therapy had no significant changes post-vaccination (Figure 3f,g). It has been found that the markers of angiogenesis (VEGF-A, PDGF-AA and PDGF-AB/BB) are elevated in hospitalized patients compared with non-critical COVID-19 infection^15^, the markers of neutrophil activation (RETN, LCN2, HGF, IL-8, G-CSF) are among the most potent discriminators of critical illness^16^, and uncontrolled activation of macrophages is responsible for acute respiratory distress syndrome (ARDS) in COVID-19 patients^17^. Our data indicate that vaccination elected detectable systemic immune modulation but does not induce a COVID-like illness in healthy individuals or even patients with cancer or autoimmune diseases on B-cell depleting therapy.

In our vaccination cohort, serum samples were collected at multiple timepoints, which are two weeks after the first dose of vaccination (Visit 1), 0~3 days before the second dose of vaccination (Visit 2), and one week after the second dose of vaccination (Visit 3). We longitudinally evaluated the protein levels at different timepoints in each group. At Visit 1, there was no significant elevation of either pro-inflammatory or T cell effector markers in healthy donors (Figure 3h,i). The peak level of these proteins was observed at Visit 2, and TNF-α, IL-15, IFN-γ dominated the elevation (Figure S8a,b), which is in line with previous reports^18^. By contrast, endotheliopathy markers peaked at Visit 3 (Figure 3j and Figure S8c), representing a late-stage response to vaccination. Again, we did not find kinetic changes in the two patient groups, and the concentration of pro-inflammatory proteins post-vaccination in Autoimmune (B cell depleted) patients was even significantly reduced when compared with pre-vaccination level (Figure 3h). Taken together, these data suggest that SARS-CoV-2 vaccination has minimal effect on the immune mediator protein production in these immunosuppressed patients.

### Natural SARS-CoV-2 infection induces a higher magnitude of the pro-inflammatory signatures compared to vaccination

Next, we asked what the protein profile difference is following natural COVID-19 infection from that induced by vaccination. To this end, we performed unsupervised clustering of 302 patient serum samples based on the concentration of the 35-protein panel, leading to the identification of five proteomically distinct subpopulations (Figure 4a). Notably, Uniform Manifold Approximation and Projection (UMAP) representation color coded by patient conditions showed that the cluster 2 subset contains mostly samples from COVID-19 infected patients (Figure 4b). The differentially expressed proteins defining each cluster indicated that samples in cluster 2 have upregulated levels of protein markers regulating inflammatory and effector T cell functions (Figure 4c). The average concentration of pro-inflammatory and T cell effector proteins was consistently higher in the two COVID+ patient groups as compared to baseline HD pre-Vax samples, whereas the significance level was reduced when comparing vaccinated groups to the baseline group (Figure 4d).

**Figure 4.**
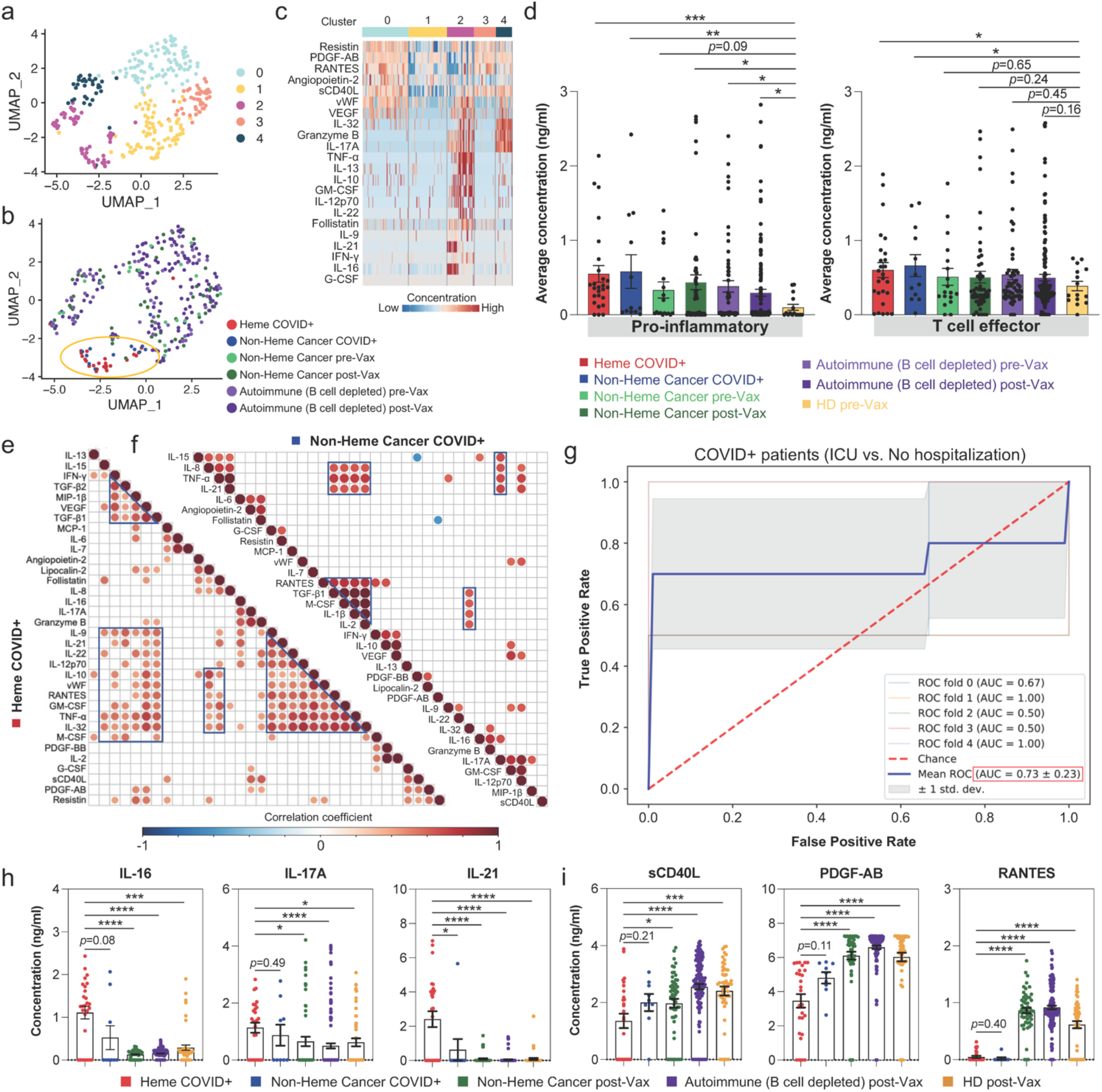
SARS-CoV-2 natural infection and vaccination induce different magnitude of pro-inflammatory signature, correlating to the level-of-care for patients. **(a)** Unsupervised clustering of 302 patient serum samples based on the concentrations of the 35-protein panel. UMAP is used to visualize the data. **(b)** UMAP representation split by patient conditions. Samples from COVID-19 infected patients are localized in cluster 2. **(c)** Differentially expressed proteins that define each cluster. **(d)** Comparisons of the average concentration of proteins regulating pro-inflammatory and T cell effector pathways between different patient/donor groups. **(e-f)** Correlation matrices of the 35-protein panel for Heme COVID+ patients (e) and Non-Heme cancer COVID+ patients (f). Only significant correlations (<0.05) are represented as dots. Pearson’s correlation coefficients from comparisons of protein concentrations across all the patients in specific groups are visualized by color intensity. Proteins were ordered by hierarchical clustering. **(g)** ROC curve for level-of-care prediction based on the pro-inflammatory protein concentrations of COVID-19 infected patients. A binomial logistic regression was used to fit the model, and a stratified fivefold cross-validation was implemented to compute the ROC and AUC. **(h-i)** Comparisons of the concentration level of specific proteins between COVID-19 infected groups and vaccinated groups. *P* values were calculated with two-tailed Mann-Whitney test. (* *P* < 0.05, ** *P* < 0.01, *** *P* < 0.001, **** *P* < 0.0001). ICU, Intensive Care Unit; ROC, receiver operator characteristic; AUC, area under the curve. UMAP, Uniform Manifold Approximation and Projection.

We next examined if SARS-CoV-2 infection conferred a different functional protein response to hematologic or non-hematologic cancer patients by interrogating the protein correlations within these two infected groups. In Heme COVID+ patients, a group of cytokines regulating inflammatory response or T cell activation, including IL-9, IL-21, IL-22, IL-12 p70, RANTES, GM-CSF, TNF-α and IL-32, were positively correlated with each other (Figure 4e). These proteins were also significantly correlated with molecules involved in T cell cytotoxicity and macrophage activation, such as IFN-γ, MIP-1β and MCP-1. While several functional T-cell cytokines were also positively correlated for Non-Heme Cancer COVID+ patients, the relationships were much weaker (Figure 4f). Prior work demonstrated that patients with severe COVID-19 had increased positive associations of proteins linked to cytokine release syndrome (CRS), which overlapped with the immune mediators identified above^19^. Thus, our data suggest that patients with hematologic cancers with COVID infection were more likely to exhibit higher disease severity than Non-Heme Cancer COVID+ patients.

The ability to predict which infected patients may become severely ill could facilitate hospital management and optimize care, which is particularly helpful for patients with cancer. Leveraging the protein data detected by our high-plex assay panel, we attempted a model to potentially predict the severity of COVID-19 infections. Patients were classified into three groups on the basis of their level-of-care requirements: admission to an Intensive Care Unit (ICU), regular hospital admission (non-ICU hospitalization) or no hospitalization. The concentrations of eight pro-inflammatory proteins or ten proteins regulating T cell effector functions were integrated into a logistic regression model using level-of-care as the predictive variable. Stratified five-fold cross-validation was used to assess the discriminative power of this model. Overall, a sensitivity of ~72% to separate ICU from no hospitalization patients with a false-positive rate of <2% was achieved using the pro-inflammatory panel [receiver operator characteristic area under the curve (ROC AUC) =0.73 ± 0.23] (Figure 4g). The sensitivity was reduced to 55% in the classification of ICU vs. non-ICU hospitalization (Figure S9a) and further reduced in the separation of non-ICU hospitalization vs. no hospitalization (Figure S9b). By contrast, the T cell effector panel could provide an improved sensitivity (~59%) to discriminate patients with non-ICU hospitalization from those without hospitalization (Figure S9c). However, this panel could not predict ICU vs. no hospitalization (Figure S9d).

To identify specific molecules that could discriminate natural infection from vaccination, we evaluated each of the proteins in our panel and compared them across groups. Among all the comparisons (Figure S10), IL-16, IL-17A, and IL-21 were found to be significantly elevated in Heme COVID+ patients compared with vaccinated groups (Figure 4h). Furthermore, the Heme COVID+ group also had significantly higher levels of the neutrophil activation marker IL-8, and the angiogenesis marker TGF-β1 than the Non-Heme Cancer COVID+ group (Figure S10), implying elevated severity in these patients with hematologic cancers^16,20^. By contrast, the concentration of the immune mediator sCD40L and PDGF-AB, a potent inducer of *CCL2* gene expression, were significantly higher in vaccinated groups (Figure 4i). Other literature has reported that these two proteins were inversely correlated with severity of COVID-19^21,22^, further suggesting increased severity signatures in infected patients. Additionally, the chemokine RANTES (CCL5) has been found to be significantly elevated from an early stage of the infection in patients with mild but not severe disease^23^. Our evaluation identified significantly higher concentration of RANTES in post-vaccination groups than infected groups, suggesting similar response characteristics between vaccination and mild infection (Figure 4i). Validation of previously identified cytokine signatures in COVID infected versus vaccinated patients further support the accuracy of our high-plex protein assay. In addition, the regression model based on above identified six proteins unambiguously separated Heme COVID+ patients from the other groups (Figure S11). Collectively, our data confirm upregulation of a group of pro-inflammatory proteins that mark higher severity of disease in patients with active COVID-19 infection, distinguish ICU admission from mild disease that does not require hospitalization, and support the observed clinical safety of vaccination compared to infection.

### A second dose vaccination boosts kinetic elevation of SARS-CoV-2 IgG bAb levels, but not in patients with B cell depletion therapy

The measurement of circulating IgG antibody binding against five protein variants from each of the spike and RBD domains in the serum samples allows us to simultaneously evaluate the bAb reactivities against wild type and several SARS-CoV-2 variants of interest. We created a “concentration heatmap” of the anti-spike and anti-RBD antibodies in each patient/donor group, showing higher levels of bAbs in patients with COVID-19 infection or groups post vaccination, except for patients with autoimmunity treated with B cell depletion therapy (Figure S12a). Next, quantitative comparisons were conducted to examine dynamic bAb changes at different timepoints after receiving vaccination. As expected, the primary vaccination effectively induced antibody responses against wild type and all the other variants in healthy donors, whereas only spike IgG bAbs showed significant increases in Non-Heme Cancer patients (Figure S12b,c). The bAb levels were sustained two weeks after the first dose of vaccination (Visit 2) (Figure 5a,b). After the secondary dose vaccination, these responses were boosted significantly in both healthy donors and Non-Heme Cancer patients, indicating the beneficial effect of the second shot. No differences were observed between the types of vaccine in this study (Figure S13a,b). In both groups antibodies effectively bounds all variants at both Visit 1 and Visit 3 (Figure 5c,d). Interestingly, we found significantly increased RBD IgG binding against B.1.1.7, B.1.351 and P.1 variants compared with the wild type protein in healthy donors while anti-spike IgG binding was uniform across variants (Figure S14a). Variant recognition by anti-spike and anti-RBD antibodies showed no differences in Non-Heme Cancer patients (Figure S14b). By contrast, both primary and secondary vaccination failed to induce robust anti-spike or RBD antibody concentrations in autoimmune patients with B cell depletion therapy (Figure 5a,b and Figure S12b,c), attributable to the elimination of circulating B-cells in this patient population. Antibody correlations were low across variants after the first dose of vaccination, especially for the RBD antibodies which improved at Visit 3 after second vaccination but not to the same levels as in the other two donor/patient groups (Figure 5e).

**Figure 5.**
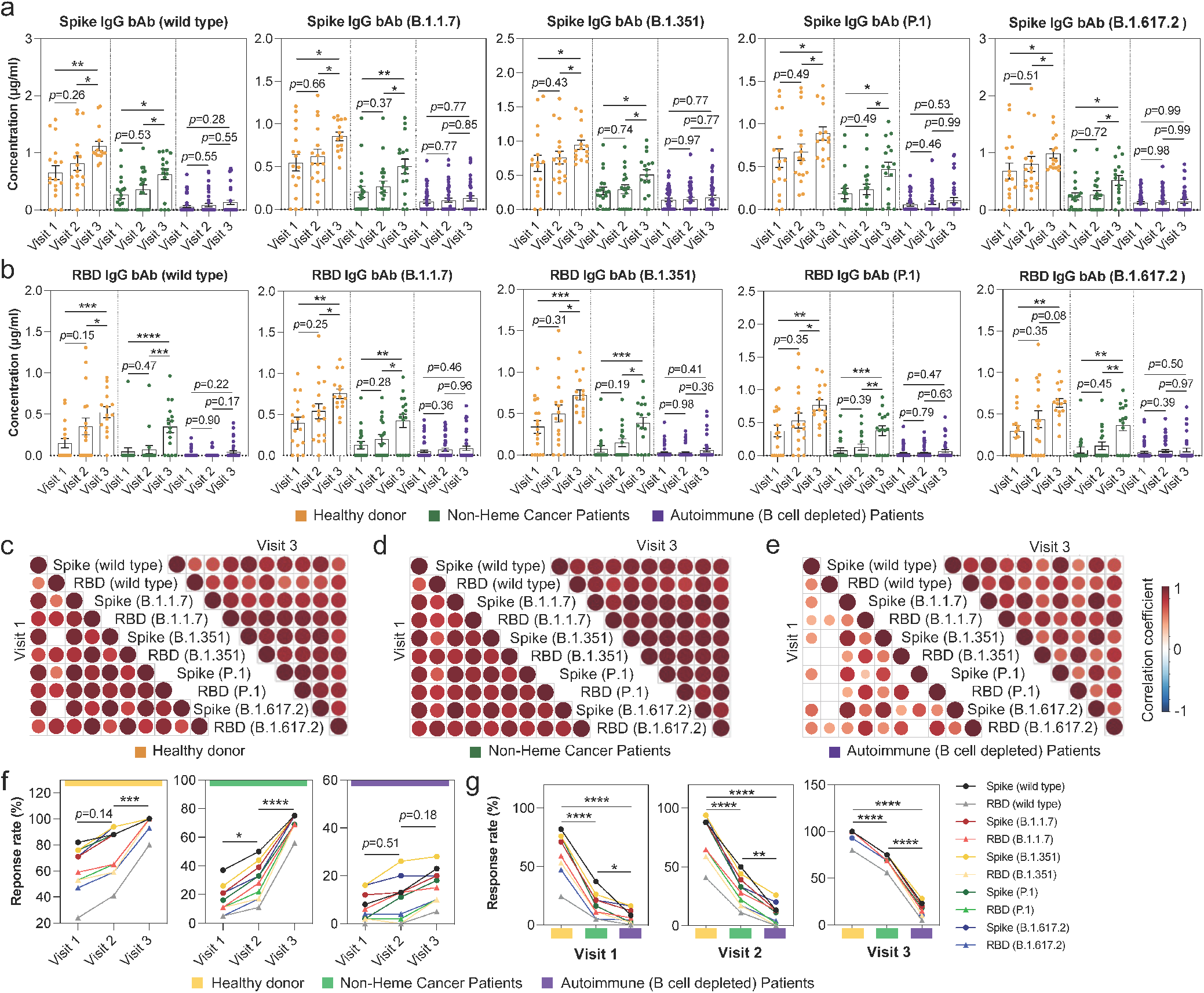
Vaccination induced circulating IgG antibody response in each patient and donor group. **(a-b)** Comparisons of the concentration of anti-spike (a) and anti-RBD (b) IgG binding against wild type or other variants between different timepoints post-vaccination. **(c-e)** Correlation matrices of the 10 SARS-CoV-2 serology panels at Visit 1 and Visit 3 in healthy donors (c); Non-Heme cancer patients (d); and autoimmune (B cell depleted) patients (e). Only significant correlations (<0.05) are represented as dots. Pearson’s correlation coefficients from comparisons of IgG antibody concentrations across all the patients in specific group are visualized by color intensity. Antibodies are listed in order of the variants’ appearance. **(f)** Comparisons of the IgG antibody response rates between different timepoints post-vaccination in each group. **(g)** Comparisons of the IgG antibody response rates between patient/donor groups at each timepoint post-vaccination. Visit 1, two weeks after the first vaccine dose; Visit 2, 0~3 days before the second vaccine dose; Visit 3, two weeks after the second vaccine dose. *P* values were calculated with two-tailed Mann-Whitney test. (* *P* < 0.05, ** *P* < 0.01, *** *P* < 0.001, **** *P* < 0.0001).

At Visit 3, we observed that anti-spike IgG levels of all the healthy individuals were above 0.5ug/ml, whereas the concentrations of several Non-Heme Cancer patients remained nearly zero, suggesting the necessity of evaluating and comparing the response rate (proportion of individuals with concentration larger than the cut-off value) between patient groups and different timepoints. The cut-off value (50 BAU/ml) of anti-spike IgG bAb provided by the World Health Organization (WHO)^24^ is equivalent to 0.3 μg/ml in our assay (Table S2), revealing protective antibody concentrations against nearly all variants in healthy donors at one week after the second dose (Visit 3) (Figure 5f). A lower response rate was noted in the Non-Heme Cancer patients at all three timepoints as compared to the healthy donor group, and the difference was further increased compared with B cell depleted patients (Figure 5g), especially for anti-RBD antibodies (Figure S15). The high-plex capacity of our device to simultaneously detect IgG antibodies and secreted proteins allowed us to examine the relationships between alterations in these functional proteins and the levels of IgG bAbs in the same serum or plasma samples. No significant correlations between protein responses and anti-spike/RBD IgG titers were detected (Figure S16). Together, these data point to a profound reduction of the second dose vaccination-induced IgG response in patients with non-hematologic cancer, and a nearly complete loss in immunocompromised patients with B cell depletion therapy.

### Hematologic malignancy patients with COVID infection exhibit impaired RBD-specific binding

We next expanded our characterization to the humoral antibody response following natural COVID-19 infection in patients with hematologic or non-hematologic cancers. As expected, the binding of SARS-CoV-2 nucleocapsid (N) protein, a RNA-binding protein critical for packaging of the viral genome^25^, was found to be substantially higher in the two infected groups as compared to vaccination groups (Figure 6a). The convalescent antibody level in Heme COVID+ patients was significantly lower than the vaccination induced antibody level in healthy donors, whereas negligible differences were found in the Non-Heme Cancer patients as compared to donors (Figure 6b). Consistently, the response rate showed a significant increase in the Non-hematologic cancer group (Figure 6c). These observations suggest that most patients with non-hematologic cancer could achieve certain levels of COVID IgG antibody seroconversion, whereas hematologic cancers and their treatment might notably impair the antibody response. No differences were observed in the comparisons between IgG titers against wild type and other variants, exclusive of enhanced binding to B.1.617.2 variant in Heme patients (Figure S17a,b). Additionally, we found significantly increased anti-RBD IgG levels in convalescent Non-Heme Cancer patients as compared to their non-infected counterparts after primary vaccination. However, the differences were no longer detectable at the later timepoints, further indicating major benefits of the second dose vaccination (Figure S18).

**Figure 6.**
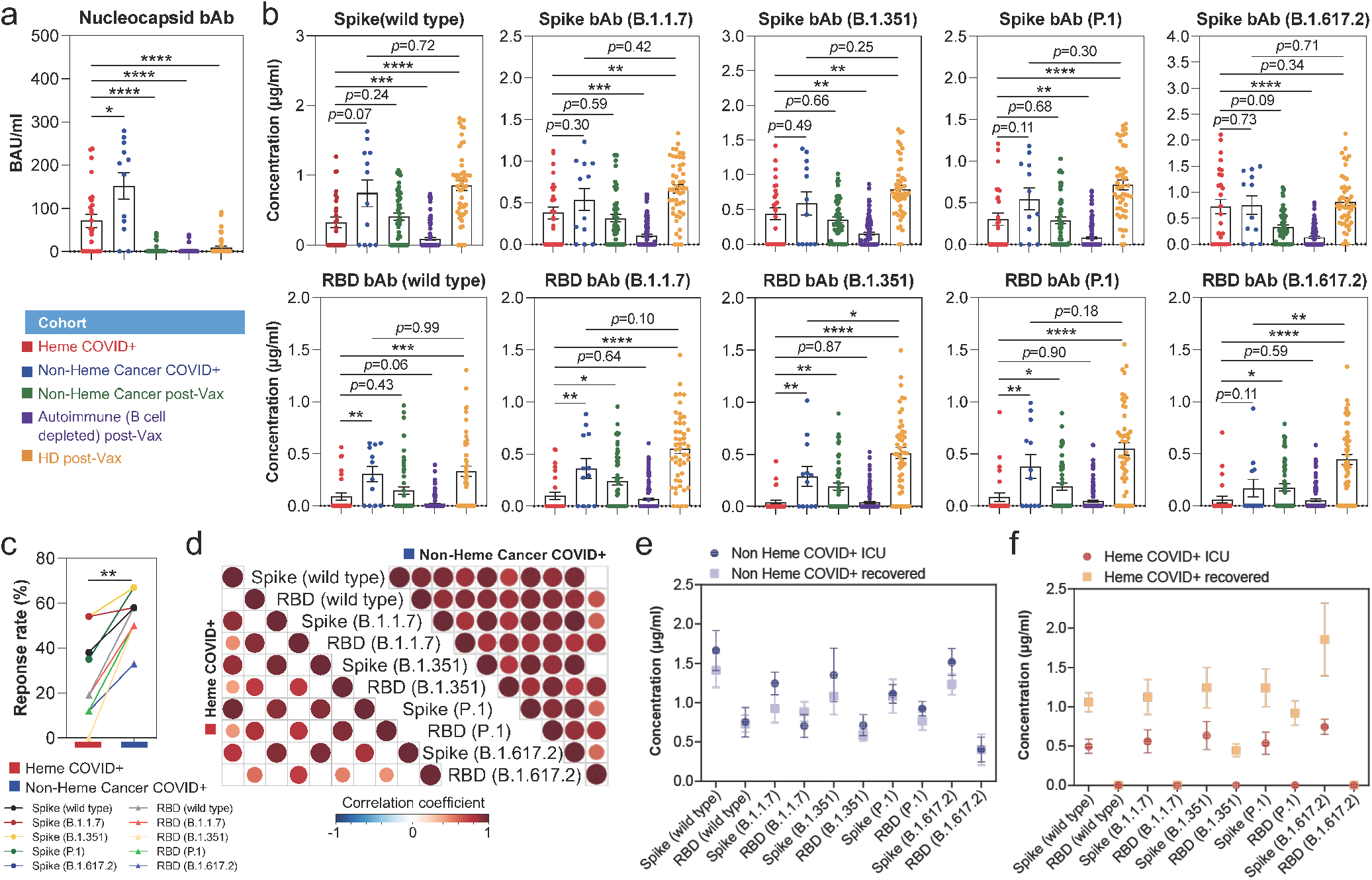
COVID-19 infected hematologic malignancy patients exhibit limited RBD-specific binding. **(a)** Comparison of the SARS-CoV-2 nucleocapsid (N) protein binding between COVID infected and post-vaccinated groups. **(b)** Comparisons of the concentration of anti-spike and anti-RBD binding against wild type and subsequent variants between COVID infected and post-vaccination groups. **(c)** Comparison of the IgG antibody response rates between COVID infected patients with hematological cancer or other types of cancer. **(d)** Correlation matrices of the 10 SARS-CoV-2 serology panels in COVID infected Heme patients and Non-Heme cancer patients. Only significant correlations (<0.05) are represented as dots. Pearson’s correlation coefficients from comparisons of IgG antibody concentrations across all the patients in specific groups are visualized by color intensity. Antibodies are listed in order of the variants’ appearance. **(e-f)** SARS-CoV-2 IgG binding antibody titers in ICU or after recovery from one Heme COVID+ (e) and one Non-Heme cancer COVID+ patient (f). *P* values were calculated with two-tailed Mann-Whitney test. (* *P* < 0.05, ** *P* < 0.01, *** *P* < 0.001, **** *P* < 0.0001). BAU, Binding Antibody Units; ICU, Intensive Care Unit.

Consistent with their reduced ability to elicit effective IgG responses, the correlations between spike and RBD binding were largely absent in COVID infected Heme patients (Figure 6d), similar to the pattern we observed in the B-cell depleted group (Figure 5e). By contrast, significantly positive associations were noted for Non-Heme Cancer patients. Again, no correlations were seen between circulating proteins and IgG antibodies in the two infected groups (Figure S19). We next longitudinally evaluated IgG bAb titers at two timepoints, in ICU or after recovery, from a hematologic cancer COVID+ (CLL) patient and a Non-Heme Cancer COVID+ patient. Similar anti-spike/RBD IgG bAb titers were observed at two timepoints for the Non-Heme cancer patient (Figure 6e), whereas the hematologic cancer patient had significantly lower anti-spike/RBD IgG bAb titers during the ICU admission with improved antibody levels at time of recovery (Figure 6f). However, this hematologic cancer patient’s RBD antibody response either during the ICU stay or at time of recovery was nearly undetectable, suggesting a lower diversity of antibody repertoire and thereby compromised humoral immune protection against COVID-19 in some hematologic cancer patients despite the presence of anti-spike IgG antibody.

### Single-cell BCR and transcriptome sequencing identifies a limited clonotype diversity and functional deficiencies in B cells from Heme COVID+ patient

The observed differences of RBD binding between COVID infected patients with hematologic or other types of cancer prompted us to further investigate the underlying immune response mechanisms. By combining single-cell RNA sequencing (scRNA-seq) and co-measurement of B cell receptor (BCR) repertoires, we characterized the heterogeneity of circulating B cells from the above evaluated hematological cancer patient and Non-Heme Cancer patient (in ICU and after recovery) (Figure 6e,f). In total, we profiled the transcriptome and BCR clonotype of 5,856 single cells from the three samples. Due to the clonal expansion and accumulation of malignant long-lived B cells in CLL patient^26^, a majority of cells (~78%; 4,526 cells) were derived from the Heme patient (Figure S20b). Integrating and clustering all the scRNA-seq data points identified 7 subpopulations and no major batch effect was observed (Figure S20a,b). High CD5 expression has been recognized as one of the signatures in leukemic B cells in CLL^27^. Consistently, we observed a notably higher expression of the encoding gene *CD5* in cells from the Heme cancer patient compared to cells from the Non-Heme cancer patient (Figure S20c). On the basis of *CD5* and *FCER2* (encoding CD23) expression, used for differential diagnosis of CLL^27^, we classified all the Heme B cells into Leukemic or Non-Malignant B cell groups (Figure S20d,e). After filtering out leukemic-B cells, 1,803 non-malignant B cells from the Heme cancer patient were included in the downstream analysis to compare with the another patient.

Consistent with our hypothesis, we found a much lower diversity of the immune repertoire in remaining non-malignant B cells from the Heme cancer patient indicating inability to develop antibody response to diverse antigens (Figure 7a). By contrast, high clonal diversity was observed in Non-Heme Cancer patient during the time both in the ICU and after recovery. The frequency of the most common clonotype was only ~0.3-0.5%, and no difference existed for the top 5 clonotypes. To further explore the association between sample source and B cell states, UMAP clustering of all the filtered B cells was performed and 5 new clusters were identified (Figure 7b and Figure S20f). A previous report identified an expansion of late-stage B cells in patients with severe symptoms as compared to the healthy controls and patients with mild symptoms^28^. We checked the same marker gene expression and found that cells in cluster 3 exhibited elevated levels of plasma cell markers (*CD27* and *CD38*)^29^, whereas cells in cluster 1 were highly enriched for *IGHD* expression, marker for naïve B-cells (Figure 7c). The cell proportion of Heme sample in cluster 1 was found to be notably lower than that of Non-Heme samples while the proportion in cluster 3 was higher (Figure 7d), suggesting more severe COVID infection signature in Heme patient. Furthermore, considering the expression levels in the three cell groups, a significantly decreased *IGHD* and increased *CD27*/*CD38* expression in Heme cells were also observed (Figure 7e).

**Figure 7.**
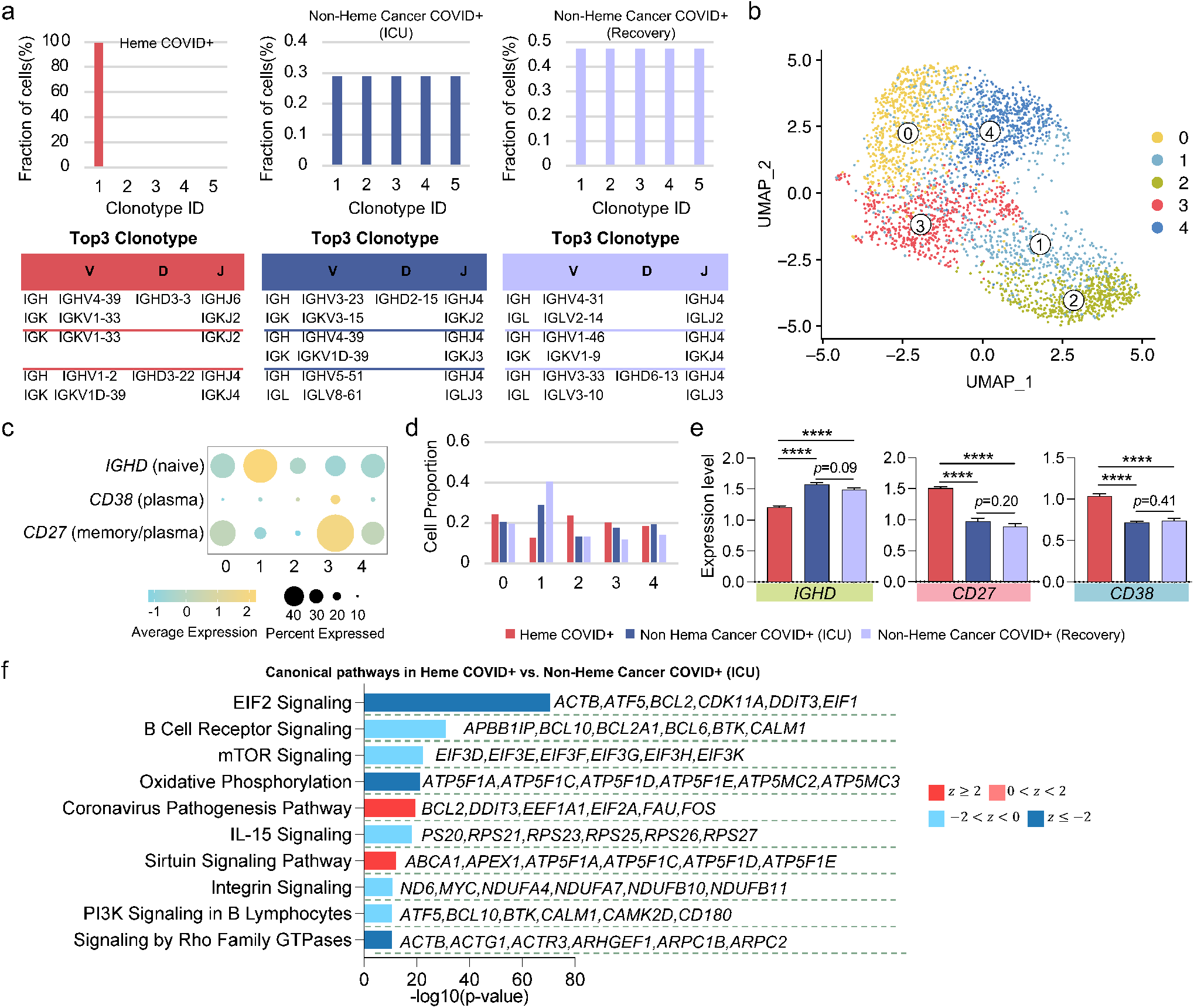
Single-cell RNA and BCR sequencing identifies low clonotype diversity and functional deficiency in B cells from the patient with hematologic cancer. (**a**) The percentage of top 5 clonotypes in each sample. (**b**) UMAP clustering of B cells from COVID infected Heme patient and Non-Heme cancer patient (two timepoints, in ICU and after recovery) identified 7 distinct subpopulations. (**c**) Dotplot of selected B cell marker gene expression in each cluster. The size of each circle represents proportion of single cells expressing the gene, and the color shade indicates normalized expression levels. (**d**) Cell proportion of each sample in each identified cluster. (**e**) Comparison of expression levels of marker genes in each sample. (**f**) Corresponding canonical pathways regulated by the highly differentially expressed genes between Heme patient vs. Non-Heme cancer patient (ICU). Pathway terms are ranked by –log 10 (*P* value). The side listed gene names represent top 6 symbolic molecular markers related to the pathway. A statistical quantity, called *z*score, is computed and used to characterize the activation level. *z*score reflects the predicted activation level (*z*<0, downregulated; *z*>0, upregulated; *z*≥2 or *z*≤−2 can be considered significant). *P* values were calculated with two-tailed Mann-Whitney test. (* *P* < 0.05, ** *P* < 0.01, *** *P* < 0.001, **** *P* < 0.0001). BCR, B cell repertoires; ICU, Intensive Care Unit. UMAP, Uniform Manifold Approximation and Projection.

To identify the unique transcriptomic signatures of each sample, we performed differentially expressed genes (DEGs) analysis among the three samples. Comparing B cells between Heme and Non-Heme (ICU) leads to the identification of top upregulated genes such as *HBB, HLA-B, NOSIP*, and top downregulated genes such as *LTB, MS4A1, CD69* (Table S3). Ingenuity Pathway Analysis (IPA)^30^ was performed to explore the biological pathways associated with the DEGs of each comparison. We found that Coronavirus pathogenesis pathway was significantly activated in B cells from Heme as compared to Non-Heme (ICU) (Figure 7f), which further confirmed that SARS-CoV-2 could induce higher severity level in immunocompromised patient. The top symbolic molecules regulating this pathway include *BCL2, DDIT3, EEF1A1, EIF2A, FAU*, and *FOS* (full gene list provided in Table S4). By contrast, a group of pathways that regulate B cell survival, proliferation, differentiation, and plasticity, including mTOR signaling, IL-15 signaling, PI3K signaling, and Signaling by Rho Family GTPases, were collectively inhibited in Heme cancer B-cells (Figure 7f). Additionally, the oxidative phosphorylation that provides metabolic support for cell activities was also found to be downregulated in Heme cancer B-cells. Similar results were obtained when comparing Heme with Non-Heme (recovery) cells (Figure S21a). Furthermore, we investigated the underlying mechanistic pathways in Non-Heme cells at ICU stage or after recovery. As expected, B cell activities and functions at ICU stage were highly upregulated in comparison with the recovery timepoint (Figure S21b). Thus, these single-cell sequencing data confirmed a markedly reduced clonotype diversity and functional deficiency in B cells from Heme COVID+ patient, which might have contributed to the observed lack of anti-RBD-specific response and likely compromised immune protection.

## Discussion

The COVID-19 outbreak has highlighted the need for easily accessible and adaptable assays to guide intervention plans, especially in high-risk groups. SARS-CoV-2 infection has particularly poor outcomes in patients with hematologic cancers and in patients with compromised B-cell immunity^31,32^. We therefore designed a portable high throughput multiplexed immune-serological assay to co-profile antibody response signature and immune mediators induced by natural infection or vaccination specifically in patients with hematologic malignancies or immunosuppressed conditions compared to patients with non-hematologic cancer or healthy donors. Our microchip assay combines high sample throughput up to 50 patient sera per chip and high degree of multiplexing for simultaneous analysis of up to a total of 50 soluble proteins and IgG antibodies. Our assay is low-cost (PDMS chips on PLL slide), fault-tolerant (5 replicates/sample), reproducible (average Pearson’s R=0.99 using 5 independent assays/devices) and characterized by high sensitivity and specificity (around 87%) using low sample volumes (8 μL) as input. It has been validated with 1,012 serum samples from patients with altered immunity and donors collected over the past two years of the pandemic.

To investigate response to infection and vaccination in immunocompromised patients versus general population and in different groups of immunocompromised patients, we analyzed 366 serum samples comprising six study cohorts: patients infected with SARS-CoV-2 with or without hematologic cancer, non-hematologic cancer patients pre- and post-vaccination, autoimmune disease patients under B cell depletion treatment pre- and post-vaccination, and healthy donors pre- and post-vaccination. For each serum sample, we simultaneously measured 35 plasma/serum proteins (divided into 6 functional categories) and SARS-CoV-2 IgG antibody against 11 different viral antigens (anti-nucleocapsid and anti-spike/RBD of wild-type virus and of other four variants). First, as expected, we noticed that the first two doses of vaccination did not induce an adequate immunological response in patients with a weakened immune system, supporting the need for additional strategies including booster doses for these patients. Specifically, we observed that immunization induced an increase in the levels and in the correlative expression of immune proteins in healthy donors, but the impact was significantly lower in non-hematologic and autoimmune patients, which is consistent with other reported studies^33^.

Then we expanded our analysis to COVID+ samples. We confirmed that natural SARS-CoV-2 infection leads to a significantly stronger pro-inflammatory signature than SARS-CoV-2 vaccination, and we demonstrated that the expression of pro-inflammatory proteins can successfully distinguish ICU admission *vs* no-hospitalization. Of note, hematologic cancer patients were unambiguously characterized by elevated levels of pro-inflammatory cytokines IL-16 and IL-17A, and of T cell effector cytokine IL-21. As previously reported^19^, high levels of these cytokines are associated with severe disease. In contrast, the concentration of sCD40L, PDGF-AB and RANTES was significantly reduced in hematologic cancer patients, further implying a higher risk of COVID-19 severity in these patients^21,22^. The evaluation of these six markers may therefore develop a multi-variant model to predict clinical outcome and to tailor treatment strategies^34^.

The analysis of anti-spike and RBD IgG bAbs supported the accuracy of our device by detecting enhanced humoral response in healthy donors one week after the second dose of COVID-19 vaccination. In contrast, as observed also for circulating immune proteins, vaccination triggered a blunted immune response in immunocompromised patients, in particular in autoimmune patients undergoing B cell depletion treatment. These data highlight the need to establish effective therapeutic strategies in autoimmune patients by balancing the benefit of booster doses, the side effect given the already dysregulated immune condition, and the administration of B cell targeting drugs^35^.

The impairment in the immune response was even more striking in COVID+ hematologic cancer patients with significantly lower levels of anti-nucleocapsid and anti-spike Abs at the time of active SARS-CoV-2 infection. Moreover, for the first time we observed that the levels of anti-RBD Abs were significantly lower in Heme COVID+ patients suggesting a limited diversity of BCR repertoire and potentially compromised immune protection even if anti-spike bAb levels were adequate. We therefore performed single-cell BCR sequencing comparing Heme *vs* Non-Heme Cancer COVID+ samples that confirmed this finding. We also observed several functional deficiencies in the immune system of these patients. Specifically, regarding non-malignant B cells in the Heme sample, we reported: i) low clonotype diversity; ii) expansion of late-stage B cells associated with severe symptoms^29,36^; and iii) enrichment in COVID-19 pathogenesis pathway and down-regulation of B cell activation pathways. Interestingly, we observed a high level of circulating IL-21 in COVID+ hematologic cancer patients, which is the signature cytokine produced by T follicular helper (Tfh) cells that promote B cell response and maintain germinal center formation and maturation in secondary lymphoid organs^37^. It seems that despite the effort to elicit Tfh response upon SARS-CoV-2 infection^38^ as characterized by elevated IL-21 level, these patients could not develop adequate B cell immunity presumably due to defective germinal center reaction, especially considering pre-existing disease-related GC dysregulation^39^.

We recognize that the current serology panel in our device does not include the latest emerging Omicron (B1.1.529) variants although these could be easily integrated in our microchip. More studies are required to define the immune response in hematologic cancer patients not only during infection but also the response to vaccination to help protect this highly vulnerable group of patients. The patient cohort needs to be enlarged to evaluate the effect of other clinical variables. Finally, we still need to examine T cell response to disentangle the complex mechanisms implicated in COVID-19 severity and immunity. Nevertheless, our newly developed high-plex assay allowed to simultaneously profile COVID-19 related plasma/serum proteins and antibodies, leading to the identification of unique signatures in immunocompromised patients, in particular hematologic cancer patients, that may be used for predicting COVID-19 prognosis and adjusting therapies for these vulnerable populations.

## Material and Methods

### Fabrication of PDMS device for high-plex immuno-serology assay

Conventional soft lithography was applied to obtain polydimethylsiloxane (PDMS) microfluidic device. A negative photoresist of SU-8 (SU-8 2025 and SU-8 2050, MicroChem) and the chrome photomasks (Range Photomasks, Lake Havasu City, AZ) was used for the molds of 25-μm-wide and 50-μm-wide microfluidic channel device. ~25 μm-thick and ~50 μm-thick SU-8 film was spin-coated on a 4-inch silicon wafer, and the molds were then fabricated following the manufacturer’s recommendations. After stir mixing GE RTV PDMS part A and part B at a 10:1 ratio, the PDMS precursor was poured onto the mold and fabricated following the degassing and curing process. The solidified PDMS slab was peeled off, and inlet and outlet holes were punched with 2 mm in diameter biopsy puncher, which can hold up to 13 μl of solution.^40^

### Sample collection

Peripheral blood was collected in red top tubes containing no anti-coagulant, allowed to clot, then centrifuged at 1500 g for 10min at 4°C within 2hrs of being drawn, the supernatant was then collected as serum. Peripheral blood samples collected in lavender Ethylenediaminetetraacetic acid (EDTA) coated tubes were layered over Ficoll and centrifuged at 1800 rpm for 20 minutes with the brake off. Plasma samples were collected from the top layer after centrifugation; peripheral blood mononuclear cells (PBMNCs) were collected from the layer below the plasma, washed with phosphate-buffered saline (PBS) counted, and then cryopreserved in dimethyl sulfoxide (DMSO) with 10% fetal bovine serum (FBS). Plasma and serum samples were stored until analysis at -80°C, and PBMNCs were stored in liquid nitrogen.

### Workflow of multiplex immuno-serology assay

A pair of PDMS device containing 50 parallel microfluidic channels was placed on the same poly-L-lysine coated slide. The first PDMS device was used for preparing the high-density barcoded array chips adapted by flow patterning method, and the second microfluidic device was employed to flow serum samples on the barcoded arrays. The channel width of the first and second PDMS microfluidic device was 25 μm and 50 μm, respectively. A 5-turn serpentine patterns was designed at the center of the first microfluidic device, and the second PDMS device had orthogonally aligned 50 parallel microfluidic channels, which can flow the loaded serum samples through the 5 replicate barcoded arrays.

For generating barcoded arrays, the first microfluidic device was placed on a clean poly-L-lysine coated slide (PLL slide, Electron Microscopy Science) and baked for 2 hours at 75°C oven to strengthen the bonding. Afterward, 4 μL of capture Ab (~4 μg/ml) or SARS-CoV-2 recombinant antigen (~0.2 mg/ml) in 1x PBS (Gibco) was loaded in each channel, and then withdrawn in using a house vacuum system for ~3 minutes. To differentiate the location of barcoded arrays in the fluorescent image, fluorescein isothiocyanate (FITC, 488 nm emission) conjugated bovine serum albumin (BSA, 1 mg/ml, Thermo Fisher Scientific) in 1x PBS was introduced to the first microfluidic channel. Following 4 hours incubation at room temperature, the capture antibodies or SARS-CoV-2 recombinant antigens were immobilized on the PLL slide. Then PLL slide with PDMS chip was soaked into the 1% BSA in 1x PBS (BSA solution) and PDMS chip was detached form the PLL slide. After removing excess capture antibodies or SARS-CoV-2 recombinant antigens with 3 times pipetting using BSA solution, the PLL slide was blocked for 1 hours at room temperature using 1 ml of BSA solution on the surface. Finally, the PLL slide was sequentially dipped into 1x PBS, 1:1 ratio of 1x PBS and deionized water (DI water), and DI water to rinse off remaining salts, and then dried using nitrogen gas. Functionalized barcoded array chips were stored in the -80 °C freezer until use (within 1 week).

To evaluate circulating proteins or SARS-CoV-2 IgG bAb in serum, the second microfluidic device was carefully aligned and attached to the first barcoded array chip. Thermal bonding was not applied to avoid degradation of immobilized panel by high temperature. A total of 8 μL serum sample from patient/donor was loaded in each channel. As similar, FITC conjugated anti-mouse IgG antibody (2 μg/ml, Cell Signaling Technology) was loaded in the first microfluidic channel to differentiate the location of loaded samples in fluorescent image as well as recognize the presence of mouse derived antibodies on PLL slide. The serum samples were withdrawn in for 20 minutes using the house vacuum system, and the flow direction was exchanged three times every 20 minutes. The circulating proteins or SARS-CoV-2 IgG Abs can be captured by the barcoded PLL slide. Afterward, the second microfluidic chip was removed from the PLL slide after soaking into BSA solution, and a cocktail of biotinylated detection antibody (~1 ng/ml per each protein panel) and phycoerythrin (PE, 532 nm emission) conjugated anti-human IgG antibody (10 ng/ml, Abcam) was loaded onto the barcoded array slide after removing excess serum using BSA solution. Following 45 minutes incubations at room temperature, allophycocyanin (APC, 635 nm emission) conjugated streptavidin (4 ng/ml, Thermo Fisher Scientific) and 1 ml of BSA solution was applied for 20 minutes and 30 min at room temperature sequentially. Finally, the fluorescent image was obtained using Genepix 4200A scanner (Molecular Devices) in 5-μm resolution after desalination and drying steps described above.

### Scanned fluorescence signal analysis

Three channels, 488 nm (FITC), 532 nm (PE) and 635 nm (APC) laser, were used to collect fluorescence signals, and each image scanning was conducted under the same power and gain values. Fluorescence intensities were evaluated with image analysis tool in Genepix Pro 6.1 software (Molecular Devices), and the mean photon counts were extracted by aligning a 20 × 20 μm^2^ square array template from not only intersection of panel and serum but also a region without microfluidic channel for serum sample. 5 square arrays from a region without second microfluidic channel was used as on-chip control to provide threshold for each panel in each replicate, and the threshold of each panel was defined as the mean value of the 5 square plus 3 times standard deviation. Only the values higher than threshold were log2 transformed after subtracting threshold, whereas values below the threshold were set as 0. If more than 2 out of 5 replicates were lower than the threshold, the data point was considered as low quality and also set as 0. Afterward, fluorescence intensity data was converted to concentration or BAU/ml unit using the titration curve shown in Figure 1d-f.

For quantitative evaluation, titration test was conducted using serial 2-fold dilution of recombinant antigens or SARS-CoV-2 Abs for the 46-plex panel, and each curve was fitted by applying hyperbola equation in a non-linear regression model using Prism 9 (GraphPad). The value of Kd and Bmax for each protein or antibody was calculated by the equation below.

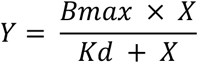

Where Bmax is the maximum number of binding sites, Kd is the ligand concentration that binds to half the receptor sites at equilibrium, respectively. Bmax and Kd values of each protein or antibody were listed in Table S5.

### Single-cell RNA sequencing and BCR sequencing

Cryopreserved PBMCs were thawed and the dead cell removal kit (Myltenyi Biotec) was applied to sort out dead cells. To enrich B cells, PBMCs were stained with anti-CD19 MicroBeads (Miltenyi Biotec) and loaded on a MACS Column that is placed in the magnetic field as per the manufacturer’s instructions. B cells from each sample were then labeled with hashtag multiplexing antibodies (BioLegend) according to the manufacture’s protocol and pooled together. Chromium Next GEM Single Cell 5’ Reagent Kits v2 (10x Genomics) were employed to perform the multi-omics profiling of enriched B cells. Briefly, stained cells were loaded onto the Chromium Next GEM Chip K, Gel Beads-in-emulsion (GEMs) were generated, and cells were lysed followed by the capture of poly-adenylated mRNA onto beads. The transcripts were reverse transcribed inside each GEM, and the barcoded full-length cDNA was PCR-amplified. Then, the amplified cDNA from mRNA and from cell surface hashtags were separated by size selection for generating V(D)J, 5ʹ gene expression libraries and hashtag libraries, respectively. Finally, the constructed libraries were sequenced on the paired-end 150bp Novaseq platform (Illumina).

### Single-cell RNA-seq data processing

The sequenced data was aligned and quantified using the CellRanger multi pipeline (version 7.0.0, 10x Genomics) against the GRCh38 human reference genome. Cells from each demultiplexed sample were first filtered based on two metrics: 1) the number of detected genes per cell must be between 200 to 5000; 2) the proportion of mitochondrial gene counts (UMIs from mitochondrial genes / total UMIs) must be less than 10%. Then, the gene expression data was normalized using Seurat sctransform.^41^ To perform batch effect correction, the Seurat v4 anchor-based integration workflow was used with the default parameter setup.^42^ Finally, the “integrated” data assay was reduced to two dimensions using UMAP for visualization, with 30 computed PCs as input. The unique cell barcodes of filtered cells were used to match and identify the BCR V(D)J gene sequences from the annotated contig files.

### Ingenuity Pathway Analysis

Ingenuity Pathway Analysis (IPA, QIAGEN) was used to understand the underlying signaling pathways.^30^ Here, DEGs distinguishing each sample, the corresponding fold change value, p value, and adjusted p value of each gene were loaded into the dataset. Ingenuity knowledge base (genes only) was used as reference set to perform Core Expression Analysis. B cell related signaling were selected from identified top canonical pathways to represent major functional profiles of each sample. The z-score was used to determine activation or inhibition level of specific pathways.

### Statistics

All the statistical analyses were performed with Prism 9 (GraphPad). Mann Whitney test was used to compare the specific observations between two groups. A *P* value<0.05 was considered statistically significant.

## Acknowledgments

The authors thank Dr. Troy Kemp from NCI Frederick National Laboratory for Cancer Research for evaluating the detection sensitivity and specificity of blinded samples. The authors also thank Dr. Lei Wang for his help with microfabrication. The microfluidic devices were fabricated in the Yale West Campus cleanroom. Computational data analysis was conducted with the Yale High Performance Computing clusters (HPC). The research was supported by Serological Science Network (SeroNet) led by National Institutes of Health (NIH) (to R.F. and S.H.; Grant number 1U01CA260507).

## Author contributions

R.F., S.H. and D.A.H. conceived the project. D.K., Z.B. and G.B. conducted the experiments. Z.B., D.K. and G.B. performed data analysis. J.V., Y.L., N.C.B., N.L., H.A., M.M.T., W.L.S. and E.E.L. helped with the collection and organization of patient samples. S.K., L.G. and J.M.K. contributed to the cohort patient consent and recruitment. M.K.R., J.E.W. and T.S.G. performed tests using Quest Diagnostics assay. Z.B., G.B., D.K., S.H. and R.F. wrote the manuscript. All authors contributed to the editing of the manuscript.

## Data and materials availability

Raw scRNA-seq data for this study are in preparation for submitting to the NCBI Gene Expression Omnibus (GEO) database (accession number pending). The other relevant data needed to evaluate the conclusions in the paper are present in the paper and/or the Supplementary Materials.

## References

1 Wang, K. et al. Memory B cell repertoire from triple vaccinees against diverse SARS-CoV-2 variants. Nature 603, 919–925, doi:10.1038/s41586-022-04466-x (2022).

2 Abdul-Jawad, S. et al. Acute Immune Signatures and Their Legacies in Severe Acute Respiratory Syndrome Coronavirus-2 Infected Cancer Patients. Cancer Cell 39, 257–275 e256, doi:10.1016/j.ccell.2021.01.001 (2021).

3 Mehta, V. et al. Case Fatality Rate of Cancer Patients with COVID-19 in a New York Hospital System. Cancer Discov 10, 935–941, doi:10.1158/2159-8290.CD-20-0516 (2020).

4 Robilotti, E. V. et al. Determinants of COVID-19 disease severity in patients with cancer. Nat Med 26, 1218–1223, doi:10.1038/s41591-020-0979-0 (2020).

5 Andrews, N. et al. Covid-19 Vaccine Effectiveness against the Omicron (B.1.1.529) Variant. New England Journal of Medicine 386, 1532–1546, doi:10.1056/NEJMoa2119451 (2022).

6 Griffiths, E. A. & Segal, B. H. Immune responses to COVID-19 vaccines in patients with cancer: Promising results and a note of caution. Cancer Cell 39, 1045–1047, doi:10.1016/j.ccell.2021.07.001 (2021).

7 Tamariz-Amador, L. E. et al. Immune biomarkers to predict SARS-CoV-2 vaccine effectiveness in patients with hematological malignancies. Blood Cancer J 11, 202, doi:10.1038/s41408-021-00594-1 (2021).

8 Amanat, F. et al. A serological assay to detect SARS-CoV-2 seroconversion in humans. Nat Med 26, 1033–1036, doi:10.1038/s41591-020-0913-5 (2020).

9 Padoan, A., Cosma, C., Sciacovelli, L., Faggian, D. & Plebani, M. Analytical performances of a chemiluminescence immunoassay for SARS-CoV-2 IgM/IgG and antibody kinetics. Clin Chem Lab Med 58, 1081–1088, doi:10.1515/cclm-2020-0443 (2020).

10 Whitman, J. D. et al. Evaluation of SARS-CoV-2 serology assays reveals a range of test performance. Nat Biotechnol 38, 1174–1183, doi:10.1038/s41587-020-0659-0 (2020).

11 Swank, Z. et al. A high-throughput microfluidic nanoimmunoassay for detecting anti-SARS-CoV-2 antibodies in serum or ultralow-volume blood samples. Proc Natl Acad Sci U S A 118, doi:10.1073/pnas.2025289118 (2021).

12 Rodriguez-Moncayo, R. et al. A high-throughput multiplexed microfluidic device for COVID-19 serology assays. Lab Chip 21, 93–104, doi:10.1039/d0lc01068e (2021).

13 Lu, Y. et al. Highly multiplexed profiling of single-cell effector functions reveals deep functional heterogeneity in response to pathogenic ligands. Proc Natl Acad Sci U S A 112, E607–615, doi:10.1073/pnas.1416756112 (2015).

14 Meizlish, M. L. et al. A neutrophil activation signature predicts critical illness and mortality in COVID-19. Blood Adv 5, 1164–1177, doi:10.1182/bloodadvances.2020003568 (2021).

15 Pine, A. B. et al. Circulating markers of angiogenesis and endotheliopathy in COVID-19. Pulmonary Circulation 10, 2045894020966547, doi:10.1177/2045894020966547 (2020).

16 Meizlish, M. L. et al. A neutrophil activation signature predicts critical illness and mortality in COVID-19. Blood Advances 5, 1164–1177, doi:10.1182/bloodadvances.2020003568 (2021).

17 Roy, R. K. et al. Macrophage Activation Syndrome and COVID 19: Impact of MAPK Driven Immune-Epigenetic Programming by SARS-Cov-2. Frontiers in immunology 12, 763313–763313, doi:10.3389/fimmu.2021.763313 (2021).

18 Bergamaschi, C. et al. Systemic IL-15, IFN-γ, and IP-10/CXCL10 signature associated with effective immune response to SARS-CoV-2 in BNT162b2 mRNA vaccine recipients. Cell Rep 36, 109504, doi:10.1016/j.celrep.2021.109504 (2021).

19 Lucas, C. et al. Longitudinal analyses reveal immunological misfiring in severe COVID-19. Nature 584, 463–469, doi:10.1038/s41586-020-2588-y (2020).

20 Ferreira-Gomes, M. et al. SARS-CoV-2 in severe COVID-19 induces a TGF-beta-dominated chronic immune response that does not target itself. Nat Commun 12, 1961, doi:10.1038/s41467-021-22210-3 (2021).

21 Zaid, Y. et al. Chemokines and eicosanoids fuel the hyperinflammation within the lungs of patients with severe COVID-19. J Allergy Clin Immunol 148, 368–380 e363, doi:10.1016/j.jaci.2021.05.032 (2021).

22 Ichiyama, T. et al. Analysis of serum soluble CD40 ligand in patients with influenza virus-associated encephalopathy. J Neurol Sci 239, 53–57, doi:10.1016/j.jns.2005.07.010 (2005).

23 Zhao, Y. et al. Longitudinal COVID-19 profiling associates IL-1RA and IL-10 with disease severity and RANTES with mild disease. JCI Insight 5, doi:10.1172/jci.insight.139834 (2020).

24 Ruetalo, N. et al. Long-Term Humoral Immune Response against SARS-CoV-2 after Natural Infection and Subsequent Vaccination According to WHO International Binding Antibody Units (BAU/mL). Viruses 13, doi:10.3390/v13122336 (2021).

25 Masters, P. S. Coronavirus genomic RNA packaging. Virology 537, 198–207, doi:10.1016/j.virol.2019.08.031 (2019).

26 Choi, M. Y., Kashyap, M. K. & Kumar, D. The chronic lymphocytic leukemia microenvironment: Beyond the B-cell receptor. Best Pract Res Clin Haematol 29, 40–53, doi:10.1016/j.beha.2016.08.007 (2016).

27 Darwiche, W., Gubler, B., Marolleau, J. P. & Ghamlouch, H. Chronic Lymphocytic Leukemia B-Cell Normal Cellular Counterpart: Clues From a Functional Perspective. Front Immunol 9, 683, doi:10.3389/fimmu.2018.00683 (2018).

28 Su, Y. et al. Multi-Omics Resolves a Sharp Disease-State Shift between Mild and Moderate COVID-19. Cell 183, 1479–1495 e1420, doi:10.1016/j.cell.2020.10.037 (2020).

29 von Borstel, A. et al. CD27(+)CD38(hi) B Cell Frequency During Remission Predicts Relapsing Disease in Granulomatosis With Polyangiitis Patients. Front Immunol 10, 2221, doi:10.3389/fimmu.2019.02221 (2019).

30 Krämer, A., Green, J., Pollard, J., Jr. & Tugendreich, S. Causal analysis approaches in Ingenuity Pathway Analysis. Bioinformatics 30, 523–530, doi:10.1093/bioinformatics/btt703 (2014).

31 Pagano, L. et al. COVID-19 infection in adult patients with hematological malignancies: a European Hematology Association Survey (EPICOVIDEHA). Journal of Hematology & Oncology 14, 168, doi:10.1186/s13045-021-01177-0 (2021).

32 Jones, J. M., Faruqi, A. J., Sullivan, J. K., Calabrese, C. & Calabrese, L. H. COVID-19 Outcomes in Patients Undergoing B Cell Depletion Therapy and Those with Humoral Immunodeficiency States: A Scoping Review. Pathog Immun 6, 76–103, doi:10.20411/pai.v6i1.435 (2021).

33 Fendler, A. et al. COVID-19 vaccines in patients with cancer: immunogenicity, efficacy and safety. Nature Reviews Clinical Oncology 19, 385–401, doi:10.1038/s41571-022-00610-8 (2022).

34 Merad, M., Blish, C. A., Sallusto, F. & Iwasaki, A. The immunology and immunopathology of COVID-19. Science 375, 1122–1127, doi:doi:10.1126/science.abm8108 (2022).

35 Lee, D. S. W., Rojas, O. L. & Gommerman, J. L. B cell depletion therapies in autoimmune disease: advances and mechanistic insights. Nature Reviews Drug Discovery 20, 179–199, doi:10.1038/s41573-020-00092-2 (2021).

36 Su, Y. et al. Heterogeneous immunological recovery trajectories revealed in post-acute COVID-19. medRxiv, 2021.2003.2019.21254004, doi:10.1101/2021.03.19.21254004 (2021).

37 Dvorscek, A. R. et al. IL-21 has a critical role in establishing germinal centers by amplifying early B cell proliferation. bioRxiv, 2022.2001.2021.476732, doi:10.1101/2022.01.21.476732 (2022).

38 Shaan Lakshmanappa, Y. et al. SARS-CoV-2 induces robust germinal center CD4 T follicular helper cell responses in rhesus macaques. Nature Communications 12, 541, doi:10.1038/s41467-020-20642-x (2021).

39 Basso, K. & Dalla-Favera, R. Germinal centres and B cell lymphomagenesis. Nature Reviews Immunology 15, 172–184, doi:10.1038/nri3814 (2015).

40 Liu, Y. et al. High-Spatial-Resolution Multi-Omics Sequencing via Deterministic Barcoding in Tissue. Cell 183, 1665–1681 e1618, doi:10.1016/j.cell.2020.10.026 (2020).

41 Hafemeister, C. & Satija, R. Normalization and variance stabilization of single-cell RNA-seq data using regularized negative binomial regression. Genome Biol 20, 296, doi:10.1186/s13059-019-1874-1 (2019).

42 Stuart, T. et al. Comprehensive Integration of Single-Cell Data. Cell 177, 1888–1902 e1821, doi:10.1016/j.cell.2019.05.031 (2019).

