## Supplemental figures and table1 2 5 for "Microfluidic immuno-serology assay revealed a limited diversity of protection against COVID-19 in patients with altered immunity"

### Content

**Figure S1.** Correlation matrices of titration values in the evaluation of SARS-CoV-2 anti-spike/ RBD IgG binding antibody against wild type or other variants.
**Figure S2.** The converted concentration or BAU based on the detected fluorescence intensity of each antibody or protein in the 5 samples.
**Figure S3.** The detected values of each SARS-CoV-2 IgG binding antibody in the evaluation of 101 preliminary blinded samples.
**Figure S4.** Concentrations of the selected proteins from samples used for the evaluation of assay accuracy and the comparison with a published report.
**Figure S5.** Concentration heatmap of 1012 serum samples measured using our assay.
**Figure S6.** Expression landscape of all the 35 proteins in different patient/donor groups.
**Figure S7.** Correlation matrices of the 35 proteins evaluated in our high-plex immuno-serology assay panel for healthy donors.
**Figure S8.** Comparisons of the concentration of each protein regulating specific immune response between pre-vaccination and different timepoints post-vaccination in healthy donors.
**Figure S9.** ROC curve for level-of-care prediction based on specific protein panel of COVID-19 infected patients.
**Figure S10.** Comparisons of the concentration level of specific proteins between COVID-19 infected groups and vaccinated groups.
**Figure S11.** ROC curve for group discrimination based on the concentrations of IL-16, IL-17A, IL-21, sCD40L, PDGF-AB, and RANTES of each sample in the specific group.
**Figure S12.** Expression landscape of all the 10 anti-spike/RBD IgG antibodies in different patient/donor groups and the comparison between pre- and post-vaccination.
**Figure S13.** Comparisons of the IgG antibody response between different types of vaccination in patient and donor groups.
**Figure S14.** Comparisons of the vaccination-induced IgG antibody response against wild type and different variants in patient and donor groups.
**Figure S15.** Comparisons of the IgG antibody response rate between anti-spike and anti-RBD binding in each group.
**Figure S16.** Correlation matrices of functional protein groups and SARS-CoV-2 serology panel at Visit 1 and Visit 3 in each patient or donor group.
**Figure S17.** Comparisons of the natural infection-induced IgG antibody response against wild type and different variants.
**Figure S18.** Comparisons of the IgG antibody response against wild type and different variants between infected patients and vaccinated patients at different timepoints.
**Figure S19.** Correlation matrices of functional protein groups and SARS-CoV-2 serology panel in COVID infected patient groups.
**Figure S20.** Filtering of leukemic B cells in patient sample with hematological cancer for downstream single-cell analysis.
**Figure S21.** Corresponding canonical pathways regulated by the highly differentially expressed genes between specific samples.
**Table S1.** Basic demographics of patients/donors evaluated in this study.
**Table S2.** The number of individuals with SARS-CoV-2 IgG antibody response larger than cut-off value in each patient/donor group.
**Table S5.** The value of Kd and Bmax for each protein or antibody calculated using hyperbola equation in a non-linear regression model.

| | | | | | | | | | | |
|---|---|---|---|---|---|---|---|---|---|---|
| Spike (wild type) | 1.00 | | | | | | | | | |
| RBD (wild type) | 1.00 | 1.00 | | | | | | | | |
| Spike (B.1.1.7) | 0.99 | 1.00 | 1.00 | | | | | | | |
| RBD (B.1.1.7) | 1.00 | 0.99 | 0.99 | 1.00 | | | | | | |
| Spike (B.1.351) | 1.00 | 1.00 | 0.99 | 0.99 | 1.00 | | | | | |
| RBD (B.1.351) | 0.99 | 0.99 | 0.98 | 0.99 | 0.99 | 1.00 | | | | |
| Spike (P.1) | 0.99 | 0.99 | 0.99 | 0.99 | 0.99 | 1.00 | 1.00 | | | |
| RBD (P.1) | 0.99 | 0.99 | 0.99 | 1.00 | 0.99 | 1.00 | 1.00 | 1.00 | | |
| Spike (B.1.617.2) | 0.99 | 0.99 | 0.99 | 0.99 | 0.99 | 1.00 | 1.00 | 1.00 | 1.00 | |
| RBD (B.1.617.2) | 0.99 | 0.99 | 0.99 | 1.00 | 0.99 | 1.00 | 1.00 | 1.00 | 1.00 | 1.00 |

**Figure S1. Correlation matrices of titration values in the evaluation of SARS-CoV-2 anti-spike/RBD IgG binding antibody against wild type or other variants.** Only significant correlations (<0.05) are represented as numbers.

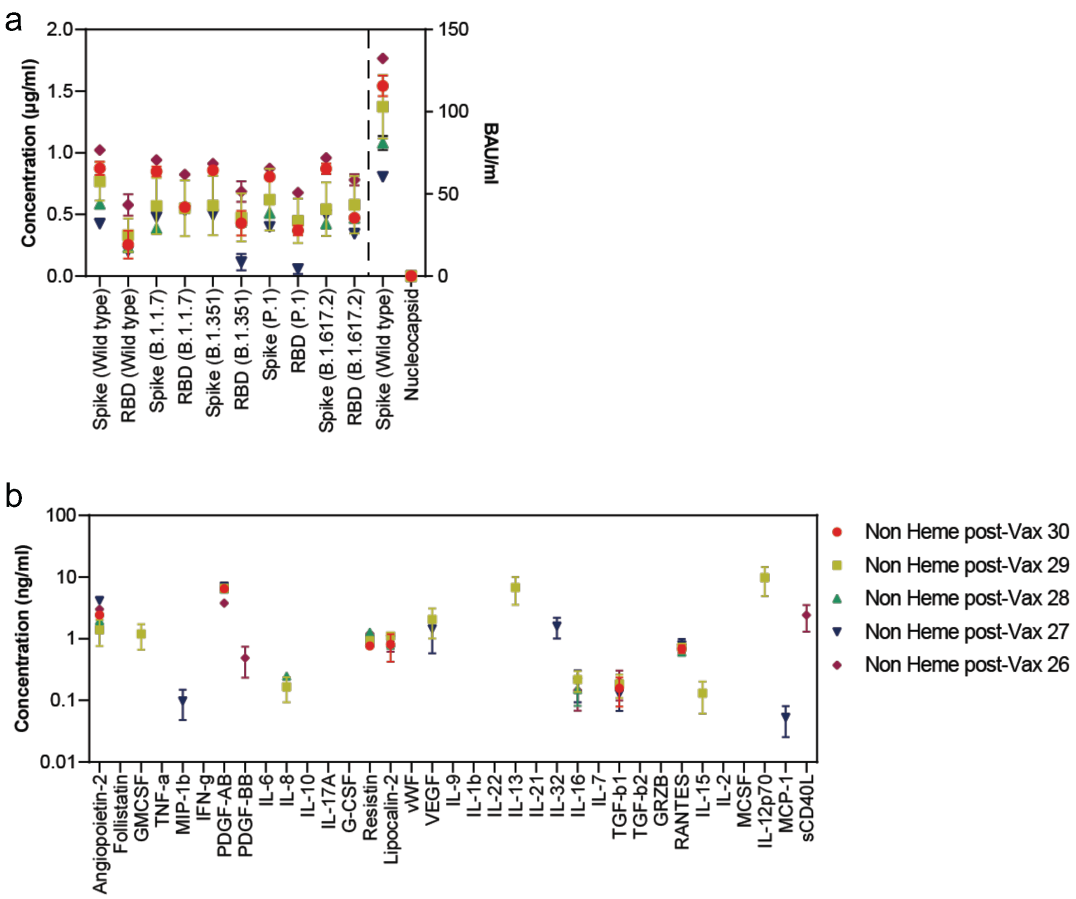


**Figure S2. The converted concentration or BAU/ml based on the detected fluorescence intensity of each antibody or protein in the 5 samples. (a)** Values of SARS-CoV-2 IgG binding antibodies. **(b)** Values of immunological proteins. Scatter plots show means ± SEM.

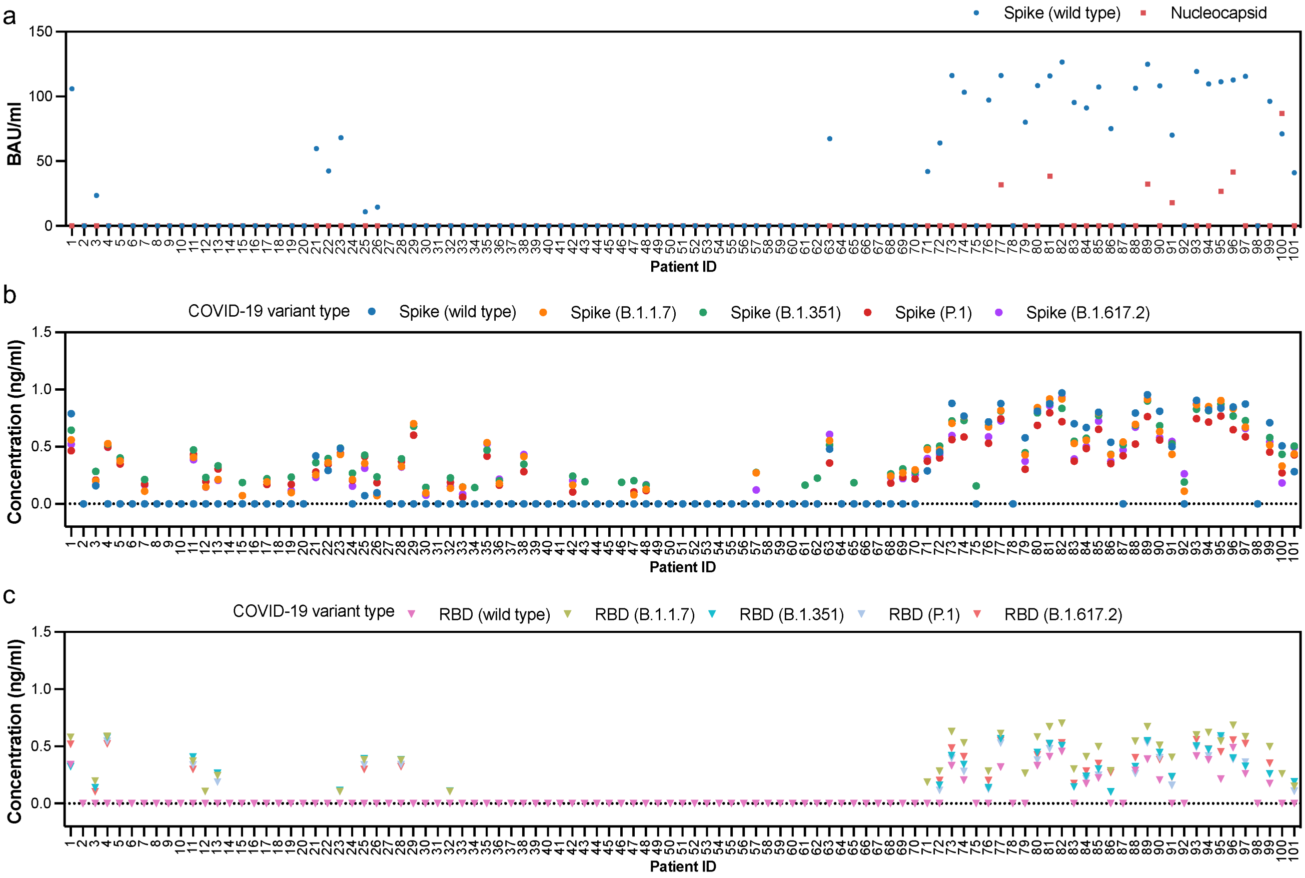


**Figure S3. The detected values of each SARS-CoV-2 IgG binding antibodies in the evaluation of 101 preliminary blinded samples**. **(a)** BAU level of wild type spike and nucleocapsid antibodies. **(b)** Concentrations of anti-spike IgG antibodies against wild type or other variants. **(c)** Concentrations of anti-RBD IgG antibodies against wild type or other variants.

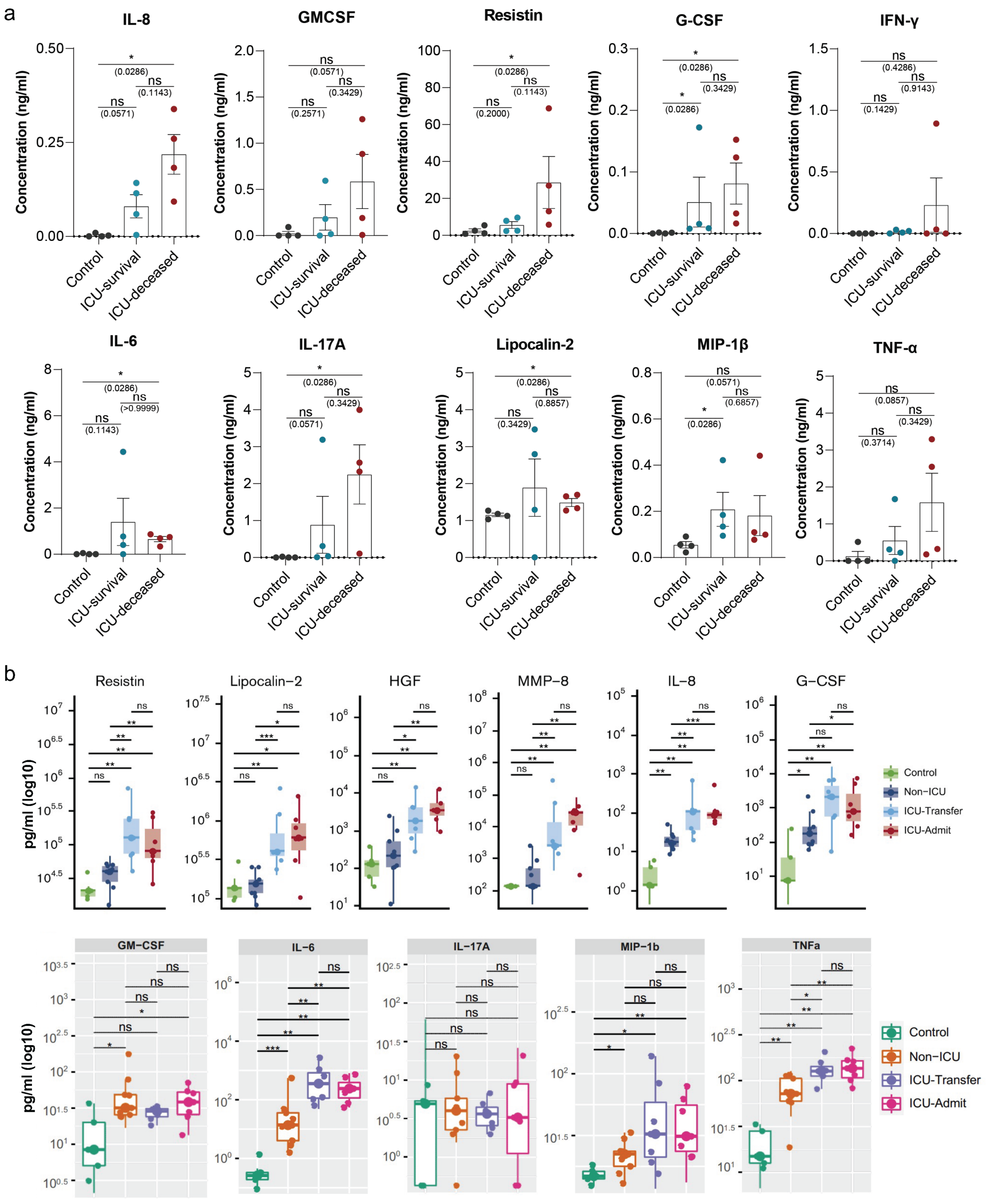


**Figure S4. Concentrations of the selected proteins from samples used for the evaluation of assay accuracy and the comparison with a published report. (a)** The results detected using our high-plex assay. Serum samples from 4 healthy controls, 4 ICU-survival patients, and 4 ICU-deceased patients were evaluated. **(b)** The results published in another report (Reference 14). Largely identical protein panel was evaluated and similar patient conditions were considered. For both our assay and published work, *P* values were calculated with two-tailed Mann-Whitney test. (* $P < 0.05$, ** $P < 0.01$, *** $P < 0.001$, **** $P < 0.0001$). ns, not significant. ICU, Intensive Care Unit; ICU-Transfer, patients later transferred to the ICU; ICU-Admit, patients directly admitted to the ICU.

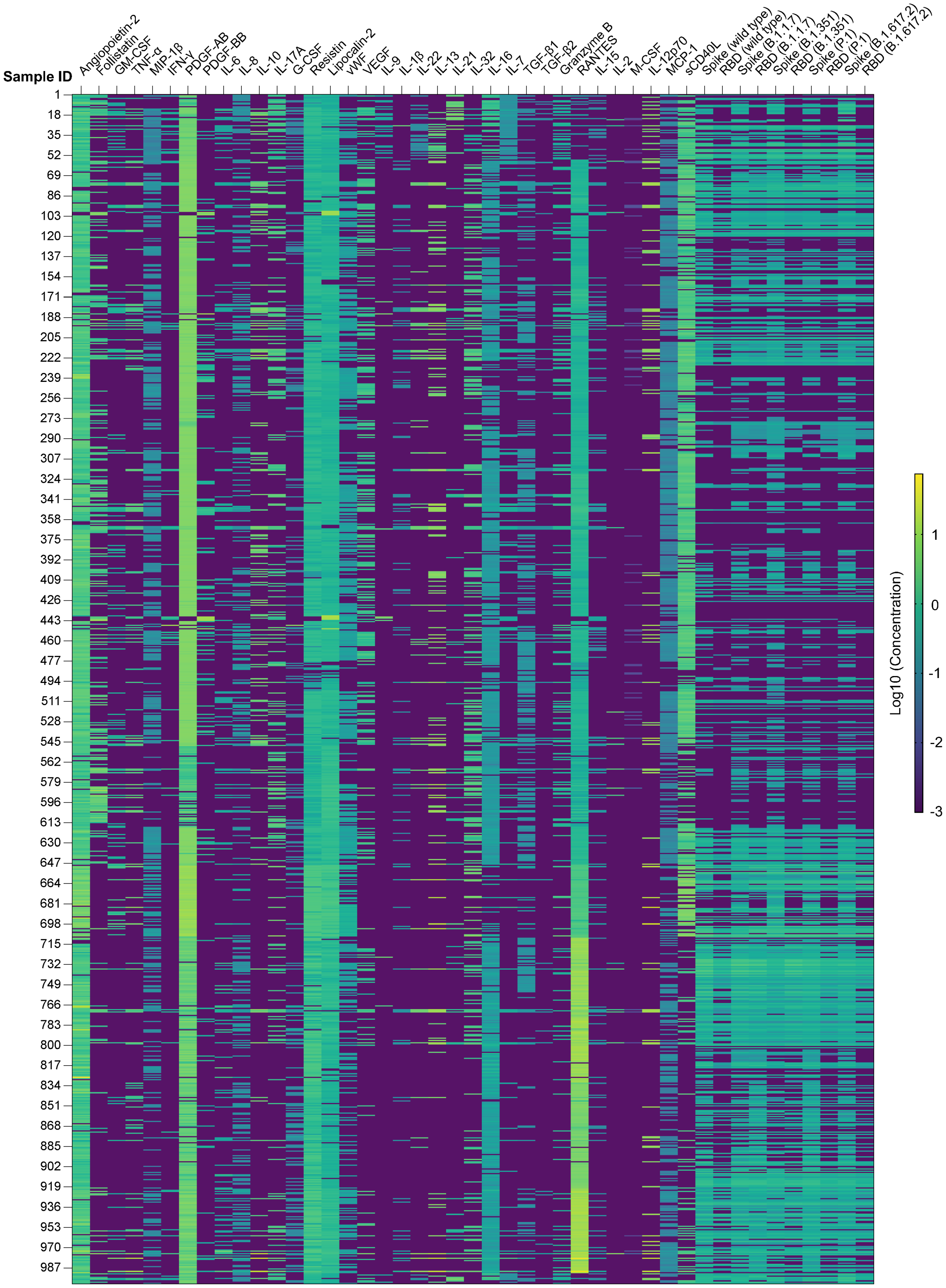


**Figure S5. Concentration heatmap of 1012 serum samples measured using our assay.** The concentration unit of protein panel is ng/ml, while that of anti-spike/RBD antibody panel is μg/ml.

a

| Sample (366 in total) | Description |
|---|---|
| Heme COVID+ (n=26) | Hematological malignancy patients with COVID19 |
| Non-Heme Cancer COVID+ (n=12) | Patients with other types of cancer and COVID19 |
| Non-Heme Cancer pre-Vax (n=18) | Patients with other types of cancer before SARS-CoV-2 vaccination |
| Non-Heme Cancer post-Vax (n=53) | Patients with other types of cancer after SARS-CoV-2 vaccination |
| Autoimmune (B cell depleted) pre-Vax (n=57) | Patients with autoimmune disease and B cell depletion therapy, before SARS-CoV-2 vaccination |
| Autoimmune (B cell depleted) post-Vax (n=136) | Patients with autoimmune disease and B cell depletion therapy, after SARS-CoV-2 vaccination |
| HD pre-Vax (n=15) | Healthy Donors before SARS-CoV-2 vaccination |
| HD post-Vax (n=49) | Healthy Donors after SARS-CoV-2 vaccination |

b

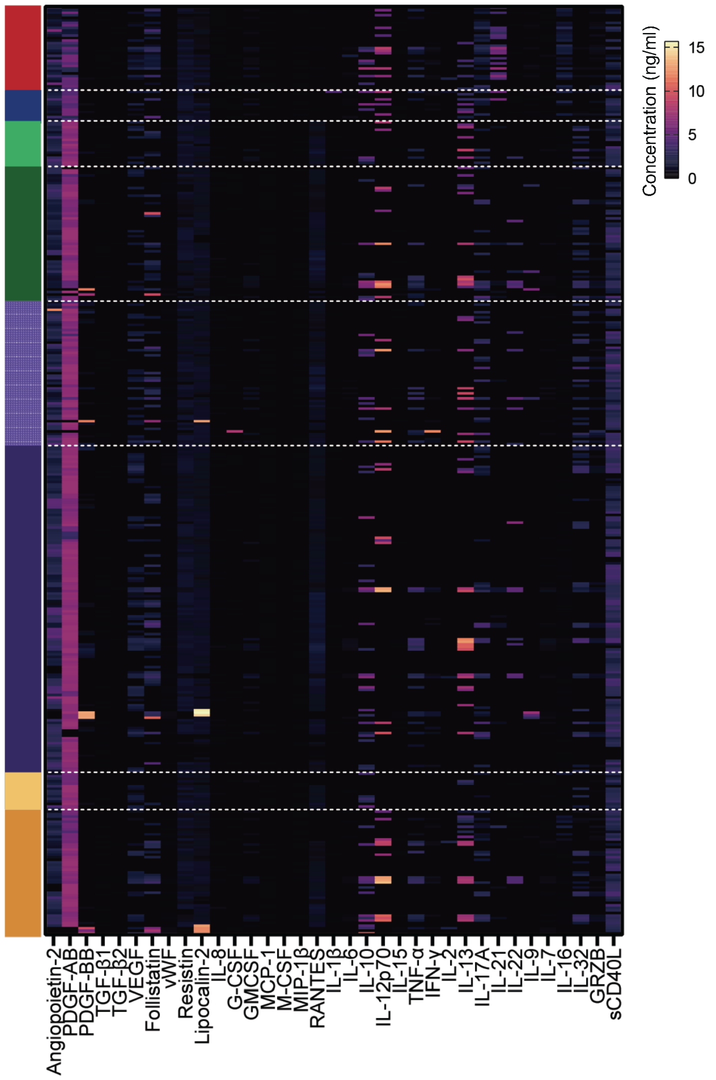


**Figure S6. Expression landscape of all the 35 proteins in different patient/donor groups. (a)** Patient or donor groups and the corresponding conditions evaluated in this work. **(b)** Concentration heatmap of 35-protein panel from all the 366 samples.

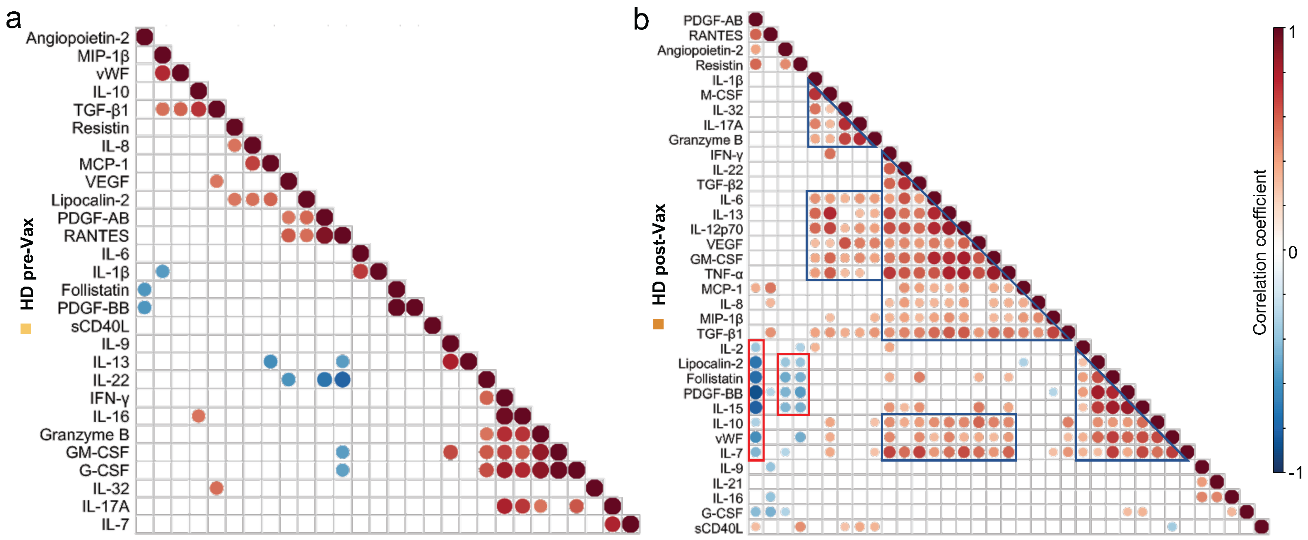


**Figure S7. Correlation matrices of the 35 proteins evaluated in our high-plex immuno-serology assay panel for healthy donors. (a)** Pre-vaccinations. **(b)** Post-vaccinations. Only significant correlations (<0.05) are represented as dots. Pearson's correlation coefficients from comparisons of protein concentrations across all the patients in specific group are visualized by color intensity. Proteins were ordered by the hierarchical clustering.

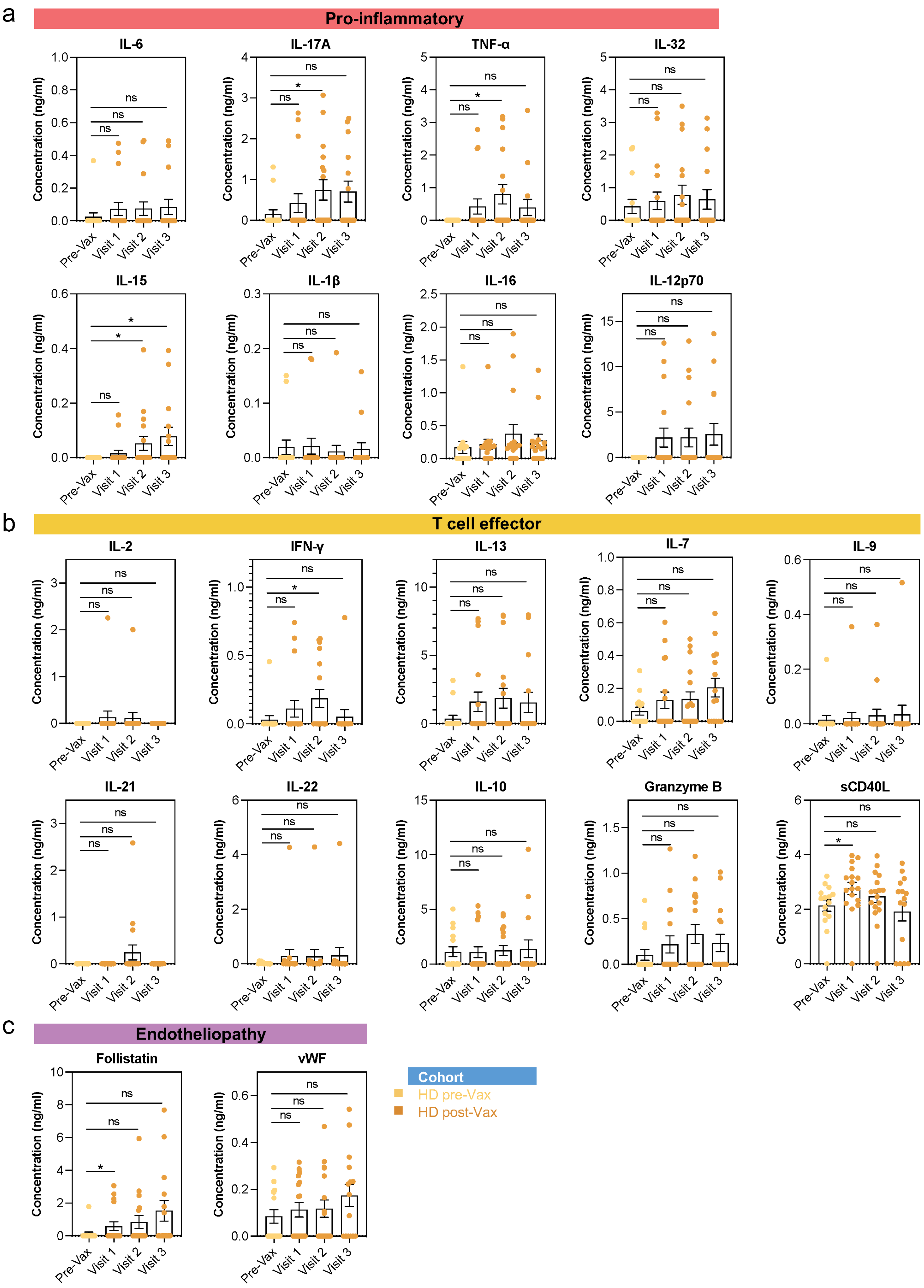

a
Pro-inflammatory
IL-6
IL-17A
TNF-α
IL-32
IL-15
IL-1β
IL-16
IL-12p70
b
T cell effector
IL-2
IFN-γ
IL-13
IL-7
IL-9
IL-21
IL-22
IL-10
Granzyme B
sCD40L
c
Endotheliopathy
Follistatin
vWF
Concentration (ng/ml)
Pre-Vax
Visit 1
Visit 2
Visit 3
ns
*
Cohort
HD pre-Vax
HD post-Vax

**Figure S8. Comparisons of the concentration of each protein regulating specific immune response between pre-vaccination and different timepoints post-vaccination in healthy donors.** (**a**) Protein panel regulating pro-inflammatory. **(b)** Protein panel regulating T cell effector **(c)** Protein panel regulating endotheliopathy**.** *P* values were calculated with two-tailed Mann-Whitney test. (* *P* < 0.05, ** *P* < 0.01, *** *P* < 0.001, **** *P* < 0.0001). ns, not significant.

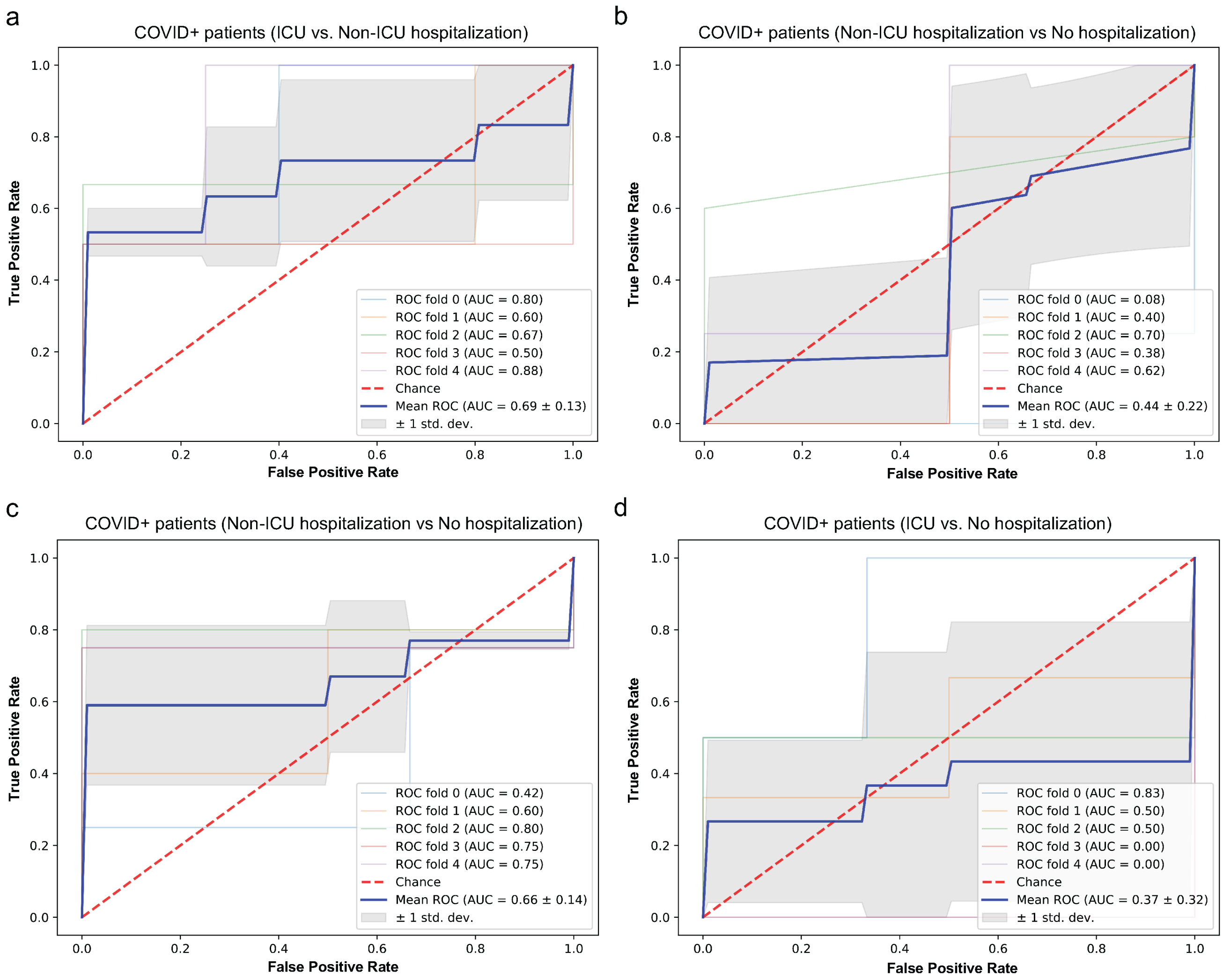


**Figure S9. ROC curve for level-of-care prediction based on specific protein panel of COVID-19 infected patients.** (**a-b**) Prediction of ICU vs. Non-ICU hospitalization (a) or Non-ICU hospitalization vs. no hospitalization (b) using protein panel regulating pro-inflammatory. **(c-d)** Prediction of Non-ICU hospitalization vs. no hospitalization (c) or ICU vs. no hospitalization (d) using protein panel regulating T cell effector. A binomial logistic regression was used to fit the model, and a stratified fivefold cross-validation was implemented to compute the ROC and AUC. ICU, Intensive Care Unit.

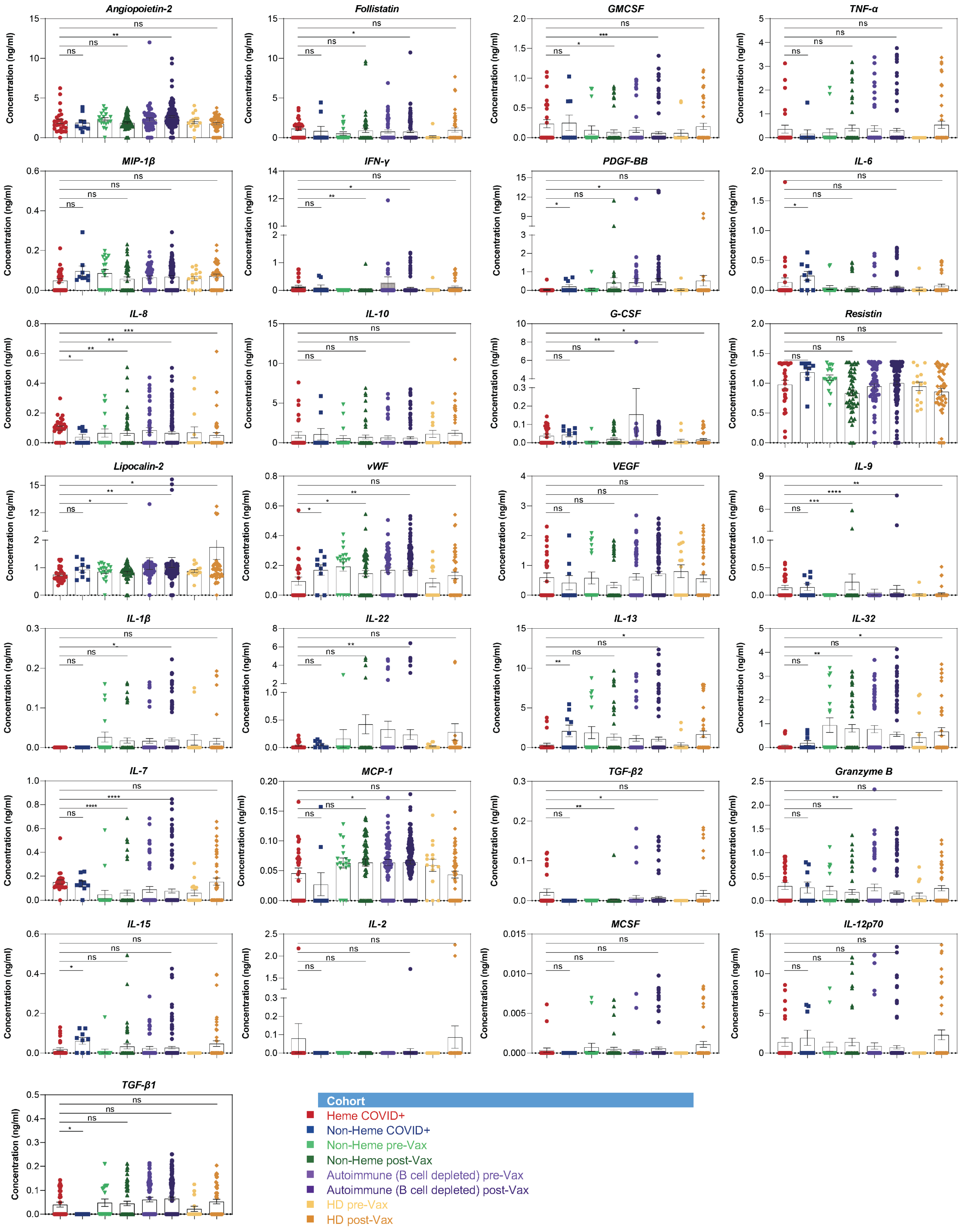


**Figure S10. Comparisons of the concentration level of specific proteins between COVID-19 infected groups and vaccinated groups.** *P* values were calculated with two-tailed Mann-Whitney test. (* *P* < 0.05, ** *P* < 0.01, *** *P* < 0.001, **** *P* < 0.0001). ns, not significant.

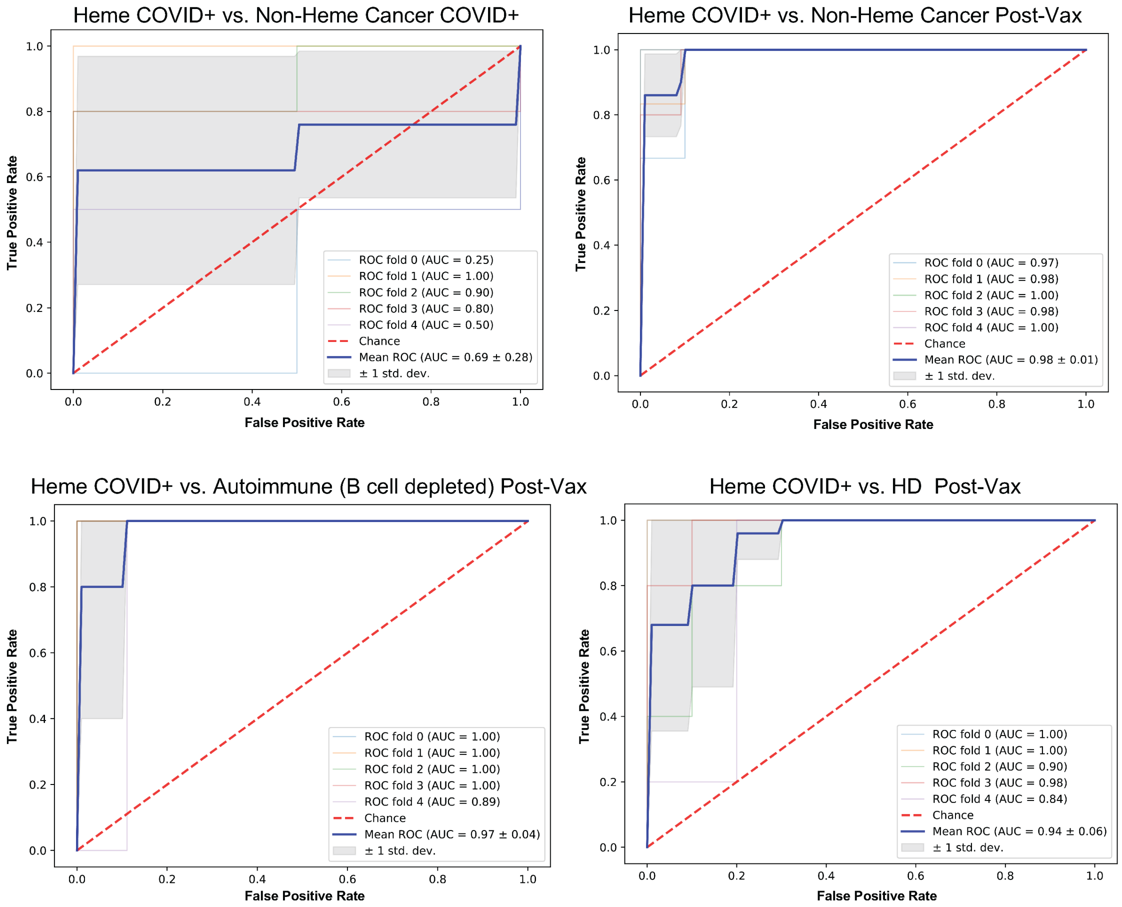


**Figure S11. ROC curve for group discrimination based on the concentrations of IL-16, IL-17A, IL-21, sCD40L, PDGF-AB, and RANTES of each sample in the specific group.** A binomial logistic regression was used to fit the model, and a stratified fivefold cross-validation was implemented to compute the ROC and AUC.

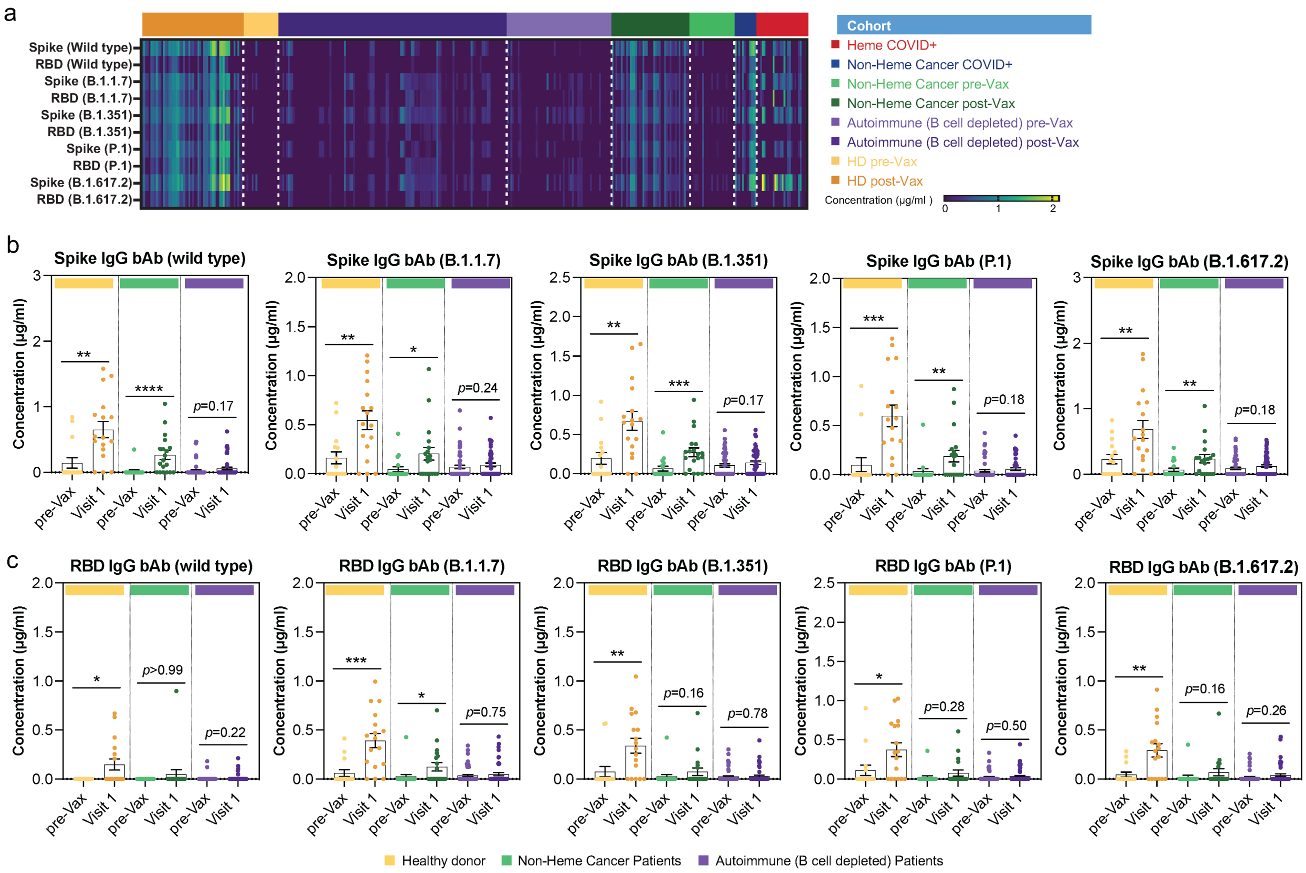


**Figure S12. Expression landscape of all the 10 anti-spike/RBD IgG antibodies in different patient/donor groups and the comparison between pre- and post-vaccination. (a)** Concentration heatmap of 10 SARS-CoV-2 IgG panel from all the 366 samples. **(b-c)** Comparisons of the concentration of anti-spike (b) and anti-RBD (c) IgG binding against wild type or other variants between pre-vaccination and post-vaccination at Visit 1. *P* values were calculated with two-tailed Mann-Whitney test. (* $P < 0.05$, ** $P < 0.01$, *** $P < 0.001$, **** $P < 0.0001$). Visit 1, two weeks after the first-dose vaccination.

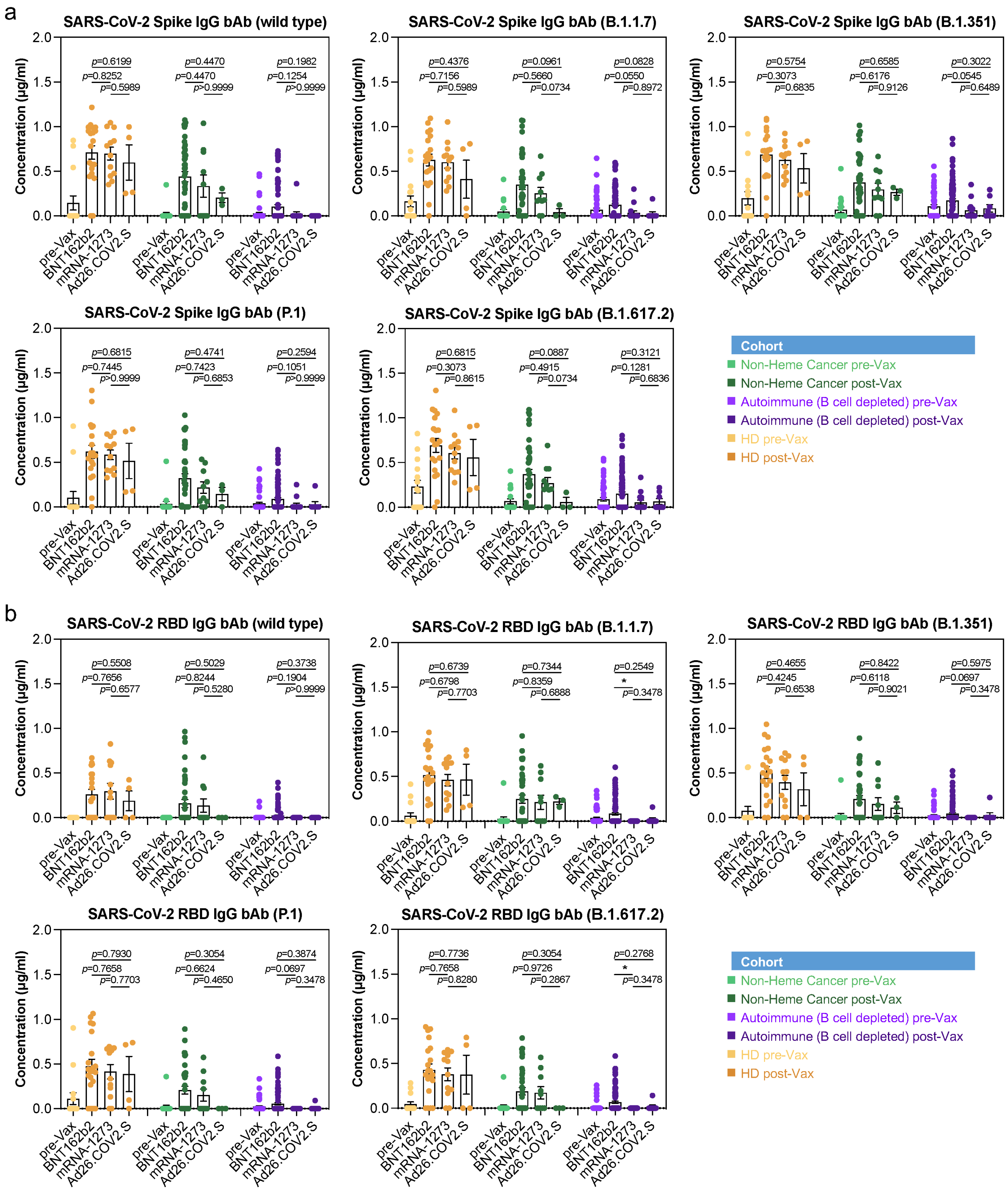


**Figure S13. Comparisons of the IgG antibody response between different types of vaccination in patient and donor groups. (a)** SARS-CoV-2 anti-spike IgG against wild type and other variants. **(b)** SARS-CoV-2 anti-RBD IgG against wild type and other variants. *P* values were calculated with two-tailed Mann-Whitney test. (* $P < 0.05$, ** $P < 0.01$, *** $P < 0.001$, **** $P < 0.0001$).

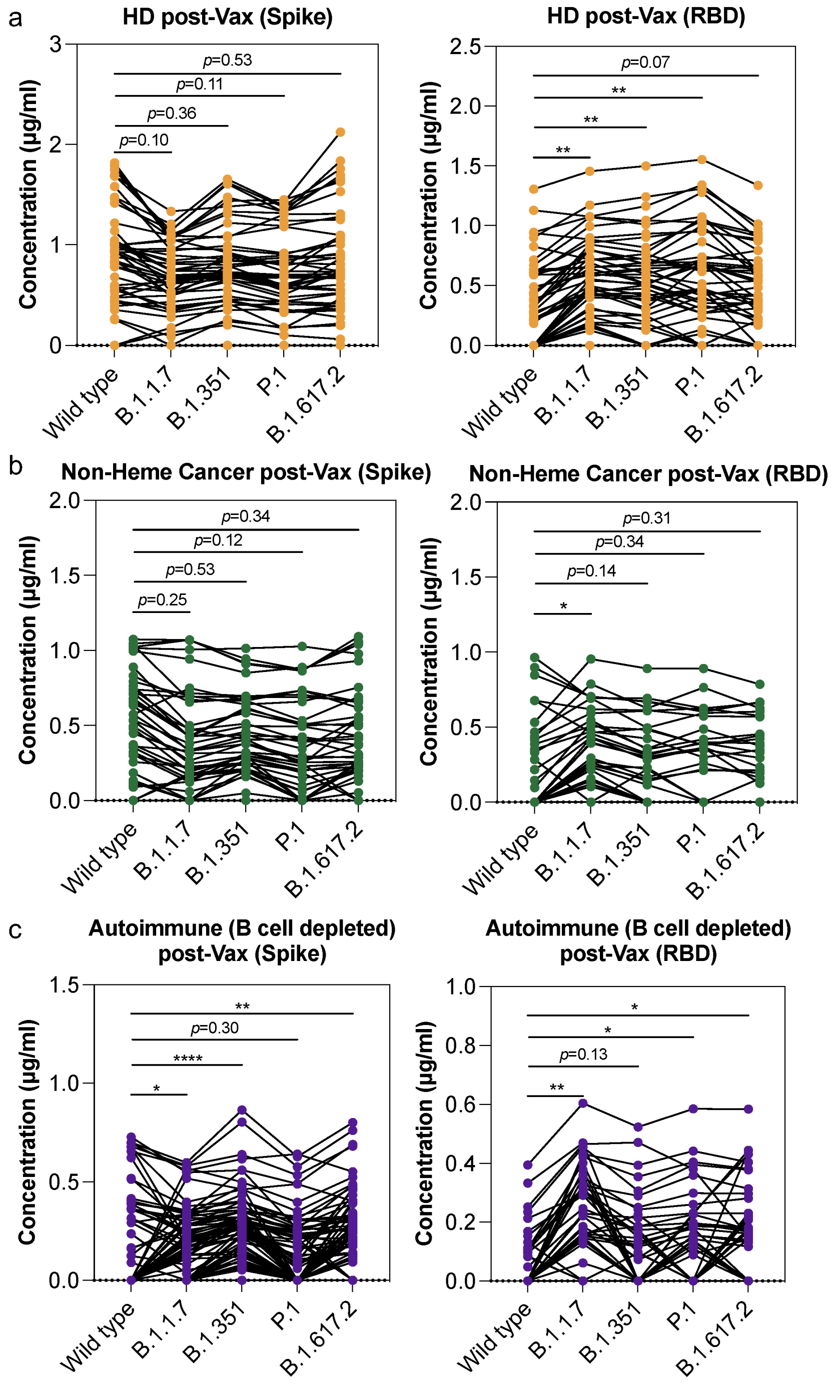


**Figure S14. Comparisons of the vaccination-induced IgG antibody response against wild type and different variants in patient and donor groups. (a)** Comparisons in healthy donor. **(b)** Comparisons in Non-Heme cancer patients. **(c)** Comparisons in autoimmune patients with B cell depletion therapy. *P* values were calculated with two-tailed Mann-Whitney test. (* $P < 0.05$, ** $P < 0.01$, *** $P < 0.001$, **** $P < 0.0001$).

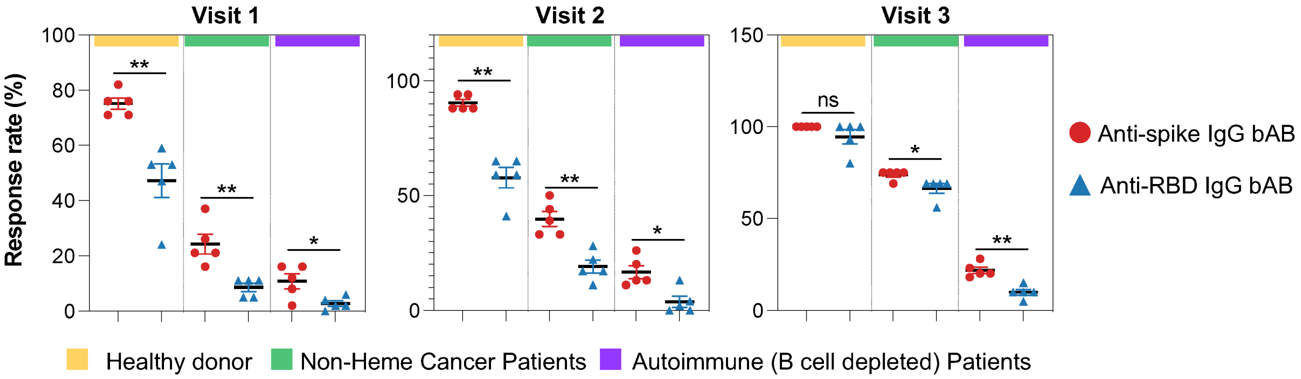


**Figure S15. Comparisons of the IgG antibody response rate between anti-spike and anti-RBD binding in each group.** *P* values were calculated with two-tailed Mann-Whitney test. (* $P < 0.05$, ** $P < 0.01$, *** $P < 0.001$, **** $P < 0.0001$). ns, not significant. Visit 1, two weeks after the first-dose vaccination; Visit 2, 0~3 days before the second-dose of vaccination; Visit 3, two weeks after the second-dose of vaccination.

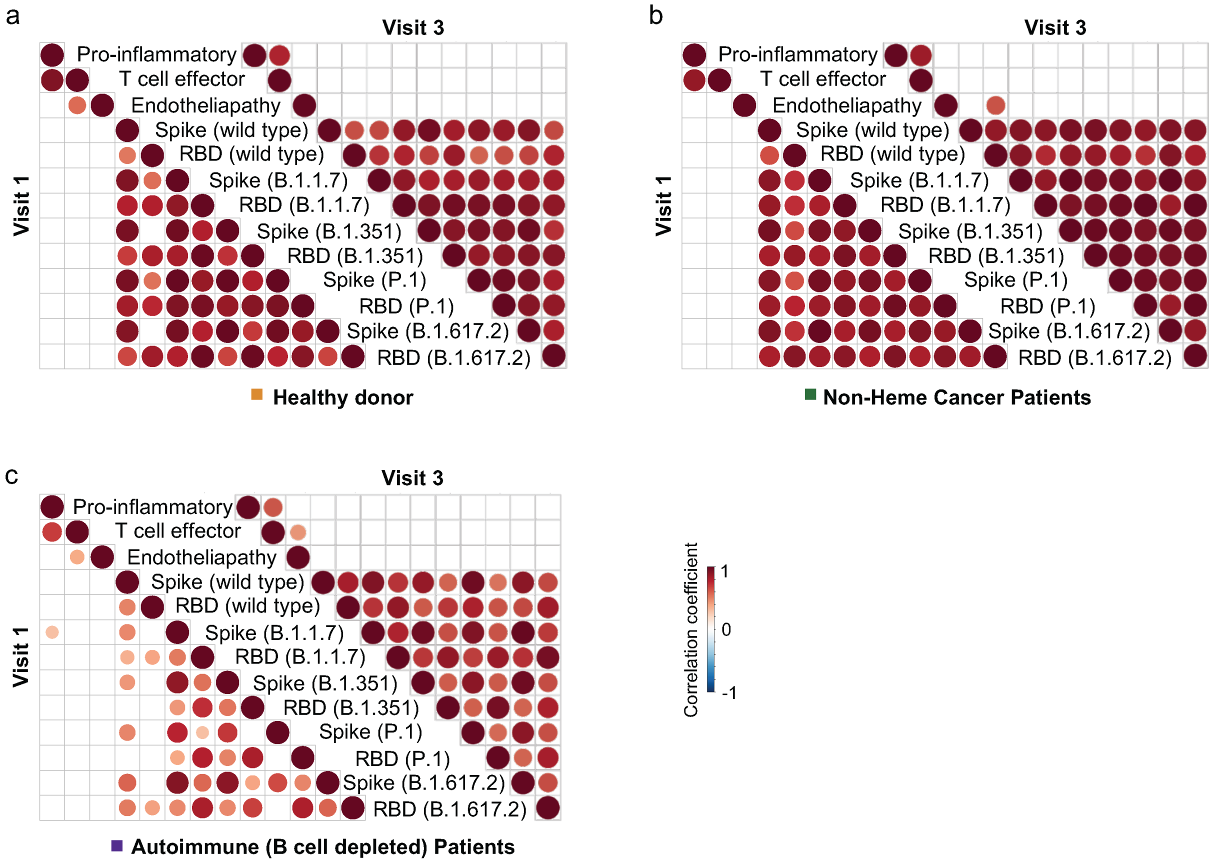


**Figure S16. Correlation matrices of functional protein groups and SARS-CoV-2 serology panel at Visit 1 and Visit 3 in each patient or donor group. (a)** Correlations in healthy donors. **(b)** Correlations in Non-Heme cancer patients. **(c)** Correlations in autoimmune patients with B cell depletion therapy. Only significant correlations (<0.05) are represented as dots. Pearson's correlation coefficients from comparisons of protein groups and IgG antibody concentrations across all the patients in specific group are visualized by color intensity. Protein groups and antibodies were listed as the original order. Visit 1, two weeks after the first-dose vaccination; Visit 3, two weeks after the second-dose of vaccination.

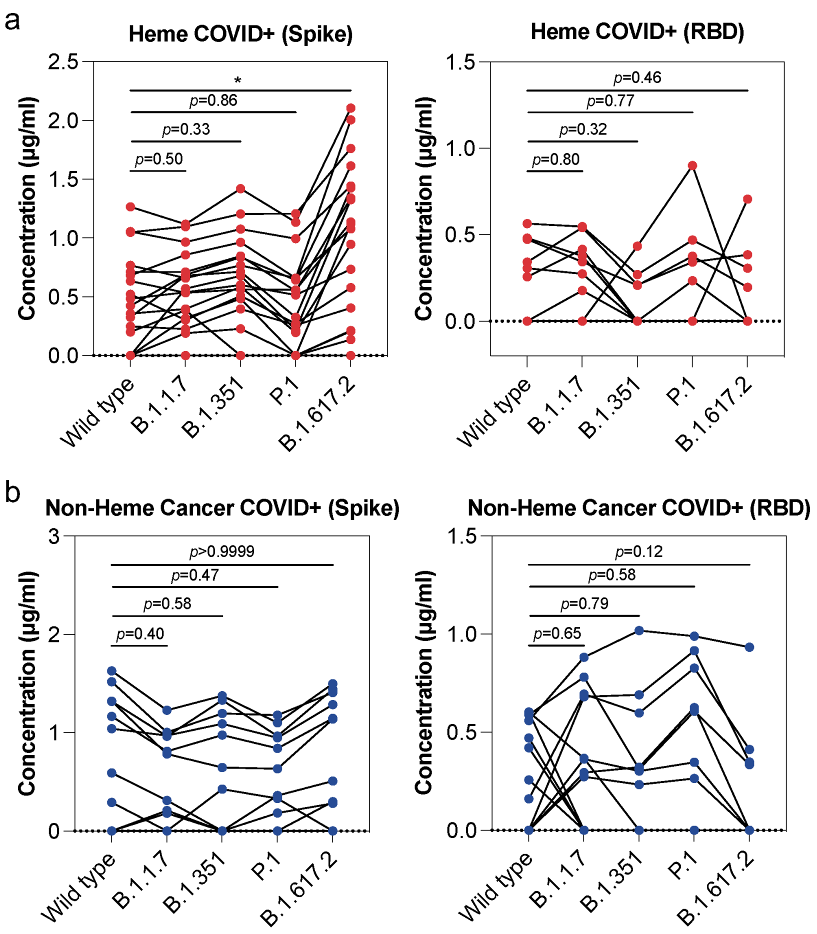


**Figure S17. Comparisons of the natural infection-induced IgG antibody response against wild type and different variants. (a)** Comparisons in Heme COVID+ patients. **(b)** Comparisons in Non-Heme cancer COVID+ patients. *P* values were calculated with two-tailed Mann-Whitney test. (* $P < 0.05$, ** $P < 0.01$, *** $P < 0.001$, **** $P < 0.0001$).

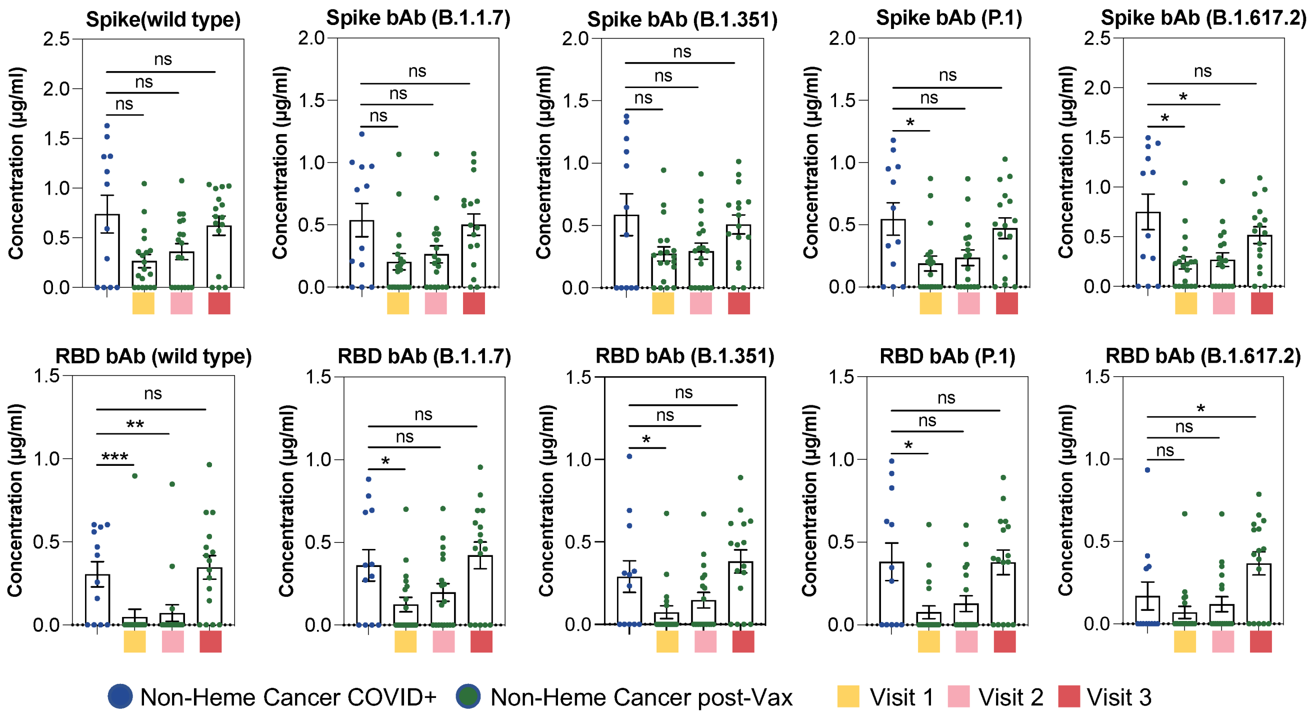


**Figure S18. Comparisons of the IgG antibody response against wild type and different variants between infected patients and vaccinated patients at different timepoints.** *P* values were calculated with two-tailed Mann-Whitney test. (* $P < 0.05$, ** $P < 0.01$, *** $P < 0.001$, **** $P < 0.0001$). ns, not significant. Visit 1, two weeks after the first-dose vaccination; Visit 2, 0~3 days before the second-dose of vaccination; Visit 3, two weeks after the second-dose of vaccination.

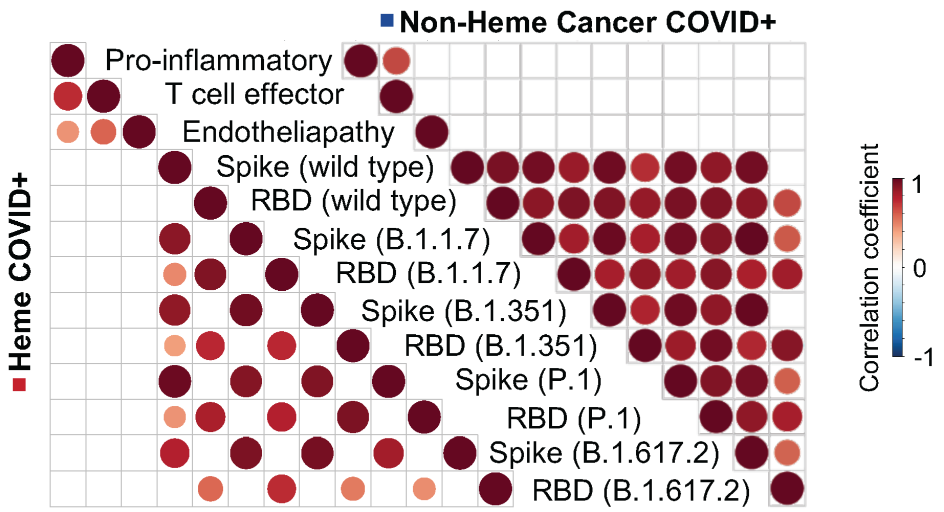


**Figure S19. Correlation matrices of functional protein groups and SARS-CoV-2 serology panel in COVID infected patient groups. (a)** Correlations in Heme COVID+ patients. **(b)** Correlations in Non-Heme cancer COVID+ patients. Only significant correlations (<0.05) are represented as dots. Pearson's correlation coefficients from comparisons of protein groups and IgG antibody concentrations across all the patients in specific group are visualized by color intensity. Protein groups and antibodies were listed as the original order.

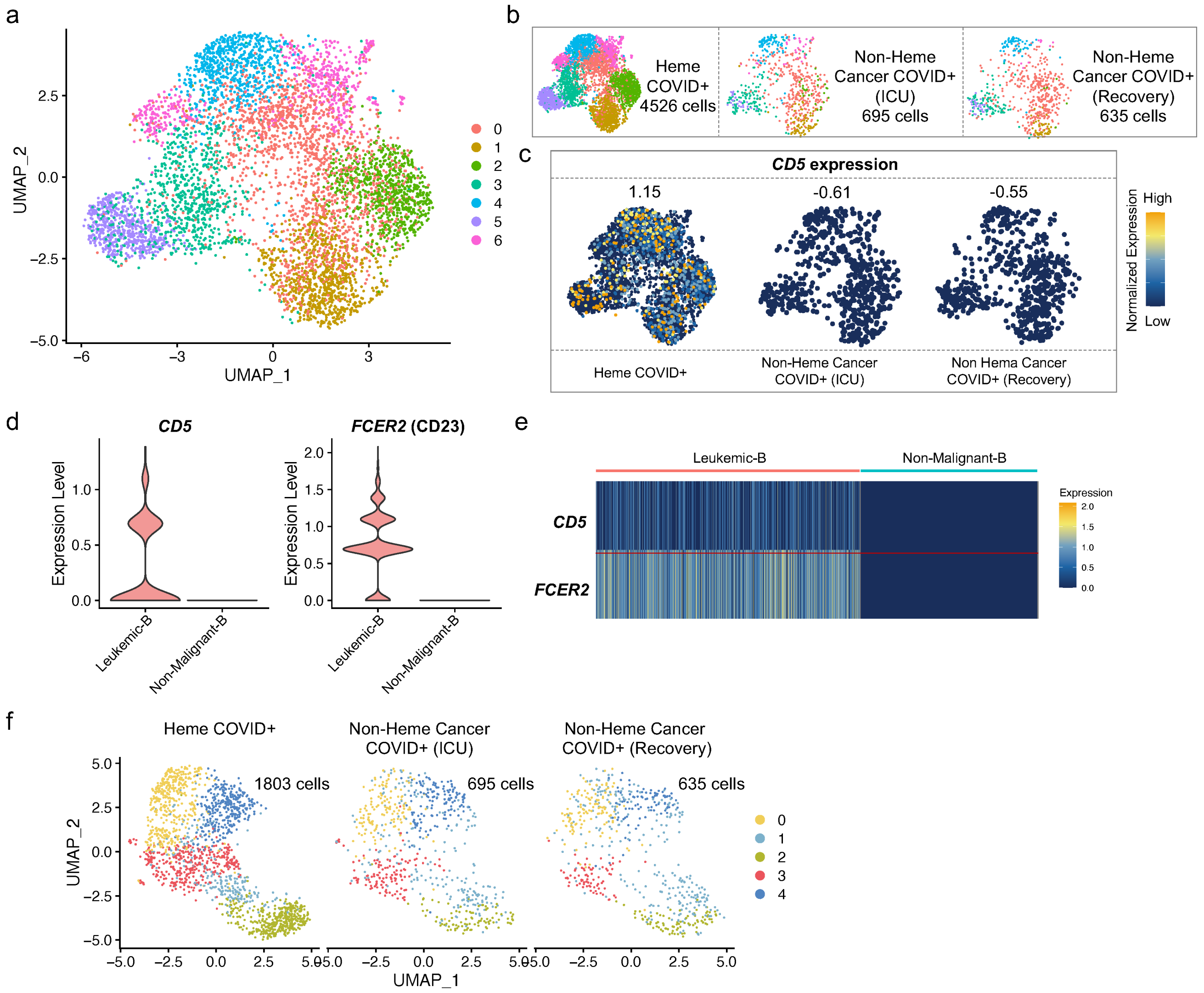


**Figure S20. Filtering of leukemic B cells in patient sample with hematological cancer for downstream single-cell analysis.** (**a**) UMAP clustering of all the B cells from COVID infected Heme patient (in ICU) and Non-Hema cancer patient (two timepoints, in ICU and after recovery). (**b**) UMAP representation split by sample resource suggests minimum batch effect. Single cell number of each sample is indicated. (**c**) The distribution of *CD5* expression split by patient identity. The number on top represents the scaled average expression of all the single cells in each sample. (**d**) Comparison of single-cell expression level of *CD5* and *FCER2* (CD23) between Leukemic-B cells and Non-Malignant-B cells. (**e**) Heatmap of *CD5* and *FCER2* (CD23) expression in Leukemic-B cells and Non-Malignant-B cells. (**f**) UMAP clustering of single cells after filtering out Leukemic-B cells, split by the patient identity. ICU, Intensive Care Unit. UMAP, Uniform Manifold Approximation and Projection.

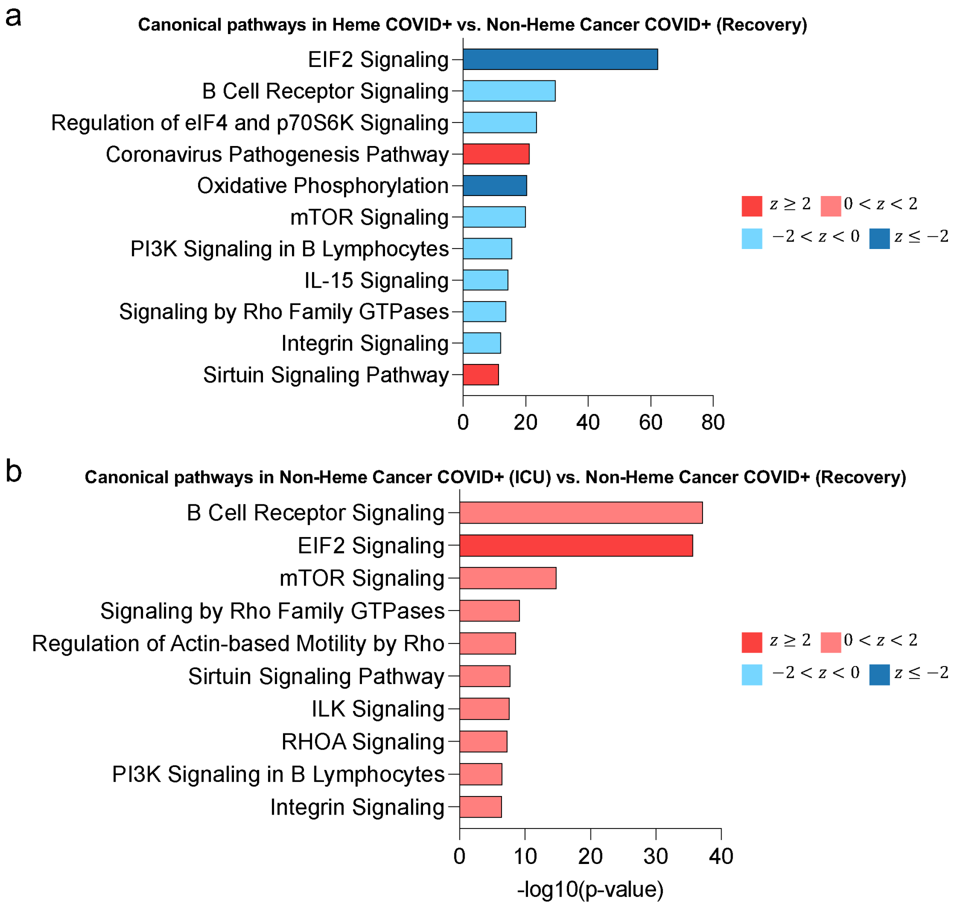


**Figure S21. Corresponding canonical pathways regulated by the highly differentially expressed genes between specific samples.** (**a**) Pathways in Heme patient (ICU) vs. Non-Heme cancer patient (recovery). (**b**) Pathways in Non-Heme cancer patient (ICU) vs. Non-Heme cancer patient (recovery). Pathway terms are ranked by –log 10 (*P* value). A statistical quantity, called $z$ score, is computed and used to characterize the activation level. $z$ score reflects the predicted activation level ($z < 0$, downregulated; $z > 0$, upregulated; $z \geq 2$ or $z \leq -2$ can be considered significant). ICU, Intensive Care Unit.

**Table S1. Basic demographics of patients/donors evaluated in this study.**

| Subject | | Heme COVID+ (n = 26) | Non-Heme Cancer COVID+ (n = 12) | Non-Heme Cancer pre- and post-Vax (n = 71) | | | | Autoimmune (B cell depleted) pre- and post-Vax (n = 193) | | | | Healthy Donor pre- and post-Vax (n = 64) | | | |
|---|---|---|---|---|---|---|---|---|---|---|---|---|---|---|---|
| | | | | Pre-Vax (n=18) | Visit 1 (n=19) | Visit 2 (n=18) | Visit 3 (n=16) | Pre-Vax (n=57) | Visit 1 (n=50) | Visit 2 (n=46) | Visit 3 (n=40) | Pre-Vax (n=15) | Visit 1 (n=17) | Visit 2 (n=17) | Visit 3 (n=15) |
| Mean age (SD), y | | 62 (15) | 57 (18) | 47 (12) | 46 (12) | 46 (12) | 47 (11) | 45 (14) | 46 (13) | 46 (12) | 47 (13) | 46 (20) | 38 (10) | 42 (18) | 43 (18) |
| Sex, n (%) | Male | 18 (69) | 5 (42) | 4 (22) | 6 (32) | 6 (33) | 4 (25) | 15 (26) | 10 (20) | 9 (20) | 8 (20) | 8 (53) | 8 (47) | 9 (53) | 7 (47) |
| | Female | 8 (31) | 7 (58) | 14 (78) | 13 (68) | 12 (67) | 12 (75) | 42 (74) | 40 (80) | 37 (80) | 32 (80) | 7 (47) | 9 (53) | 8 (47) | 8 (53) |

n represents the number of samples from each specific patient/donor group.

**Table S2. The number of individuals with SARS-CoV-2 IgG antibody response larger than cut-off value in each patient/donor group.**

| Subject | Heme COVID+ (n = 26) | Non-Heme Cancer COVID+ (n = 12) | Non-Heme Cancer pre- and post-Vax (n = 71) | | | | Autoimmune (B cell depleted) pre- and post-Vax (n = 193) | | | | Healthy Donor pre- and post-Vax (n = 64) | | | |
|---|---|---|---|---|---|---|---|---|---|---|---|---|---|---|
| | | | Pre-Vax (n=18) | Visit 1 (n=19) | Visit 2 (n=18) | Visit 3 (n=16) | Pre-Vax (n=57) | Visit 1 (n=50) | Visit 2 (n=46) | Visit 3 (n=40) | Pre-Vax (n=15) | Visit 1 (n=17) | Visit 2 (n=17) | Visit 3 (n=15) |
| *Response rate of SARS-CoV-2 IgG bAb (> 0.3 μg/ml or > 50 BAU/ml), n (%)* | | | | | | | | | | | | | | |
| Spike (wild type), BAU/ml | 10 (38) | 7 (58) | 1 (6) | 7 (37) | 9 (50) | 12 (75) | 2 (4) | 4 (8) | 6 (13) | 9 (23) | 3 (20) | 14 (82) | 15 (88) | 15 (100) |
| RBD (wild type), μg/ml | 5 (19) | 7 (58) | 0 (0) | 1 (5) | 2 (11) | 9 (56) | 2 (4) | 0 (0) | 0 (0) | 2 (5) | 0 (0) | 4 (24) | 7 (41) | 12 (80) |
| Spike (B.1.1.7), μg/ml | 14 (54) | 7 (58) | 4 (22) | 4 (21) | 7 (39) | 12 (75) | 16 (28) | 6 (12) | 6 (13) | 8 (20) | 7 (47) | 12 (71) | 15 (88) | 15 (100) |
| RBD (B.1.1.7), μg/ml | 5 (19) | 6 (50) | 1 (6) | 2 (11) | 5 (28) | 11 (69) | 11 (19) | 3 (6) | 6 (13) | 6 (15) | 3 (20) | 10 (59) | 11 (65) | 15 (100) |
| Spike (B.1.351), μg/ml | 14 (54) | 8 (67) | 6 (33) | 5 (26) | 8 (44) | 12 (75) | 23 (40) | 8 (16) | 12 (26) | 11 (28) | 7 (47) | 13 (76) | 16 (94) | 15 (100) |
| RBD (B.1.351), μg/ml | 0 (0) | 6 (50) | 1 (6) | 2 (11) | 3 (17) | 11 (69) | 8 (14) | 1 (2) | 0 (0) | 4 (10) | 2 (13) | 9 (53) | 10 (59) | 15 (100) |
| Spike (P.1), μg/ml | 9 (35) | 8 (67) | 2 (11) | 3 (16) | 6 (33) | 11 (69) | 9 (16) | 1 (2) | 5 (11) | 7 (18) | 2 (13) | 13 (76) | 15 (88) | 15 (100) |
| RBD (P.1), μg/ml | 3 (12) | 6 (50) | 1 (6) | 2 (11) | 4 (22) | 11 (69) | 8 (14) | 1 (2) | 1 (2) | 4 (10) | 4 (27) | 9 (53) | 11 (65) | 15 (100) |
| Spike (B.1.617.2), μg/ml | 14 (54) | 7 (58) | 5 (28) | 4 (21) | 6 (33) | 12 (75) | 18 (32) | 8 (16) | 9 (20) | 8 (20) | 8 (53) | 12 (71) | 16 (94) | 15 (100) |
| RBD (B.1.617.2), μg/ml | 3 (12) | 4 (33) | 1 (6) | 1 (5) | 3 (17) | 11 (69) | 6 (11) | 2 (4) | 2 (4) | 4 (10) | 3 (20) | 8 (47) | 10 (59) | 14 (93) |
| Nucleocapsid, BAU/ml | 13 (50) | 10 (83) | 0 (0) | 0 (0) | 0 (0) | 0 (0) | 0 (0) | 0 (0) | 0 (0) | 0 (0) | 3 (20) | 0 (0) | 2 (12) | 1 (7) |

n represents the number of samples from each specific patient/donor group.

**Table S5. The value of Kd and Bmax for each protein or antibody calculated using hyperbola equation in a non-linear regression model.**

| Panel | $B_{max}$ | $K_d$ | Panel | $B_{max}$ | $K_d$ |
|---|---|---|---|---|---|
| Angiopoietin-2 | 14.13 | 0.4902 | IL-1b | 12.17 | 0.05796 |
| Follistatin | 13.09 | 1.082 | IL-22 | 15.75 | 2.859 |
| GMCSF | 12.79 | 0.3752 | IL-13 | 16.84 | 5.301 |
| TNF-a | 12.93 | 1.026 | IL-21 | 14.13 | 0.5607 |
| MIP-1b | 13.41 | 0.04363 | IL-32 | 13.96 | 1.037 |
| IFN-g | 15.4 | 0.5326 | IL-16 | 14.31 | 0.0896 |
| PDGF-AB | 19.57 | 1.312 | IL-7 | 13.49 | 0.2333 |
| PDGF-BB | 15.28 | 0.4169 | TGF-b1 | 13.75 | 0.068 |
| IL-6 | 13.23 | 0.202 | TGF-b2 | 16.5 | 0.1103 |
| IL-8 | 12.7 | 0.05464 | GRZB | 14.41 | 0.4943 |
| IL-10 | 12.09 | 1.952 | RANTES | 12.72 | 0.1119 |
| IL-17A | 13.74 | 1.115 | IL-15 | 13.73 | 0.06644 |
| G-CSF | 14.13 | 0.05208 | IL-2 | 12.36 | 0.9377 |
| Resistin | 15.86 | 0.0702 | MCSF | 14.45 | 0.004836 |
| Lipocalin-2 | 12.69 | 0.2125 | IL-12p70 | 18.73 | 8.724 |
| vWF | 13.59 | 0.1284 | MCP-1 | 12.93 | 0.02396 |
| VEGF | 13.89 | 0.9005 | CD40L | 12.93 | 1.026 |
| IL-9 | 12.12 | 2.245 | SARS-CoV-2 IgG Ab (μg/ml) | 10.55 | 0.00039 |
| Spike-IgG (BAU/ml) | 10.92 | 0.06131 | Nucleocapsid-IgG (BAU/ml) | 11.29 | 0.02593 |
